# Deep conservation of head direction circuits in bees, ants and flies

**DOI:** 10.64898/2026.07.26.740564

**Authors:** Marcel E. Sayre, Valentin Gillet, Nils Ceberg, Griffin Badalamente, Atticus Pinzon-Rodriguez, Nina Griggs, Laia Serratosa Capdevila, Ruairí J.V. Roberts, Ebba Sundberg Gunnarsson, Felicia Szadaj, Saroja Ellendula, Anna Honkanen, Ajay Narendra, Stanley Heinze

## Abstract

To navigate, animal brains must continuously estimate the body’s heading in space and compare it with internal goals to guide movement. In the fruit fly *Drosophila melanogaster* a neural circuit in the central complex of the brain serves as an internal compass, exploiting a network architecture called a ring attractor^1, 2^. While this region is highly conserved and involved in navigation in many insects^3, 4^, it remains unclear whether the fly circuit represents a general blueprint for head direction computation, or whether different ecologies have driven distinct circuit solutions. Using synaptic-resolution circuit mapping, we identified homologous head direction networks in bees and ants and compared them to the fly circuit. We show that the insect head direction network is conserved across at least 300 million years of evolution. At the level of cell types and projection patterns, all studied species share a nearly identical neural layout, both qualitatively and quantitatively. At the synaptic level, however, the fly and bee circuits differed fundamentally. The distinct wiring principles of homologous neurons expose highly evolvable elements within this otherwise stable circuit. Using these differences in circuit architecture to constrain computational models, we show that both the bee and fly circuits can effectively function as ring attractors with similar, yet distinct, properties. These results demonstrate that complex neural circuits can remain stable over many hundreds of millions of years, while still offering access points for evolution to flexibly adjust neural computations to changing ecological demands, illustrating how evolution balances stability and flexibility in brain circuits.

---

Animal navigation relies on internal spatial representations in the brain. These representations vary in function and complexity, from cognitive maps in rodents^5, 6^, bats^7^, and humans^8^ to vector representations in fruit flies^9, 10^ and bees^11^. A principal feature required by any navigator is a continuous internal representation of its current heading. This is neurally implemented by head direction cells, which were initially discovered in rats^12^, but have since been found in evolutionarily distant animals, including other mammals^13, 14^, birds^15^, fish^16, 17^, and insects^18–22^. Theoretical models proposed that head direction signals could be computed by circuits arranged in a network structure known as a ring attractor^23, 24^. In this ring-like arrangement of neurons, neural signals are shaped by local excitation and long-range inhibition to generate a “bump” of confined activity that corresponds to the angular orientation of the animal and which persists even in the absence of sensory input. A physical implementation of this circuit was structurally and functionally confirmed in the fruit fly, *Drosophila melanogaster* ^1, 2, 25^. While ring attractor-like dynamics have also been identified in zebrafish^17^ and rodents^26^, mechanistic understanding of head direction coding is currently limited to the fruit fly^27^. It is unknown whether the fruit fly head direction circuit is representative of other species.

The fly head direction cells are located in the central complex^28, 29^, a brain region composed of the ellipsoid body, the protocerebral bridge, the fan-shaped body and the noduli (Figure 1a,b). The central complex is characterized by a highly regular internal organization consisting of eight repeating computational units, called columns. These columns evenly map 360°of azimuthal space around the fly, tightly coupling structure to function. This is achieved by a population of columnar cells (EPG neurons) that form an activity bump whose location within the central complex columns correlates with the fly’s heading. The position of this bump follows the angular movements of the fly, much like a biological compass needle^1, 2, 19, 30, 31^. This encoding scheme is enabled by an interplay of several recurrent neural loops, generated by anatomically defined cell types, which together form a biological ring attractor^1^.

**Figure 1:**
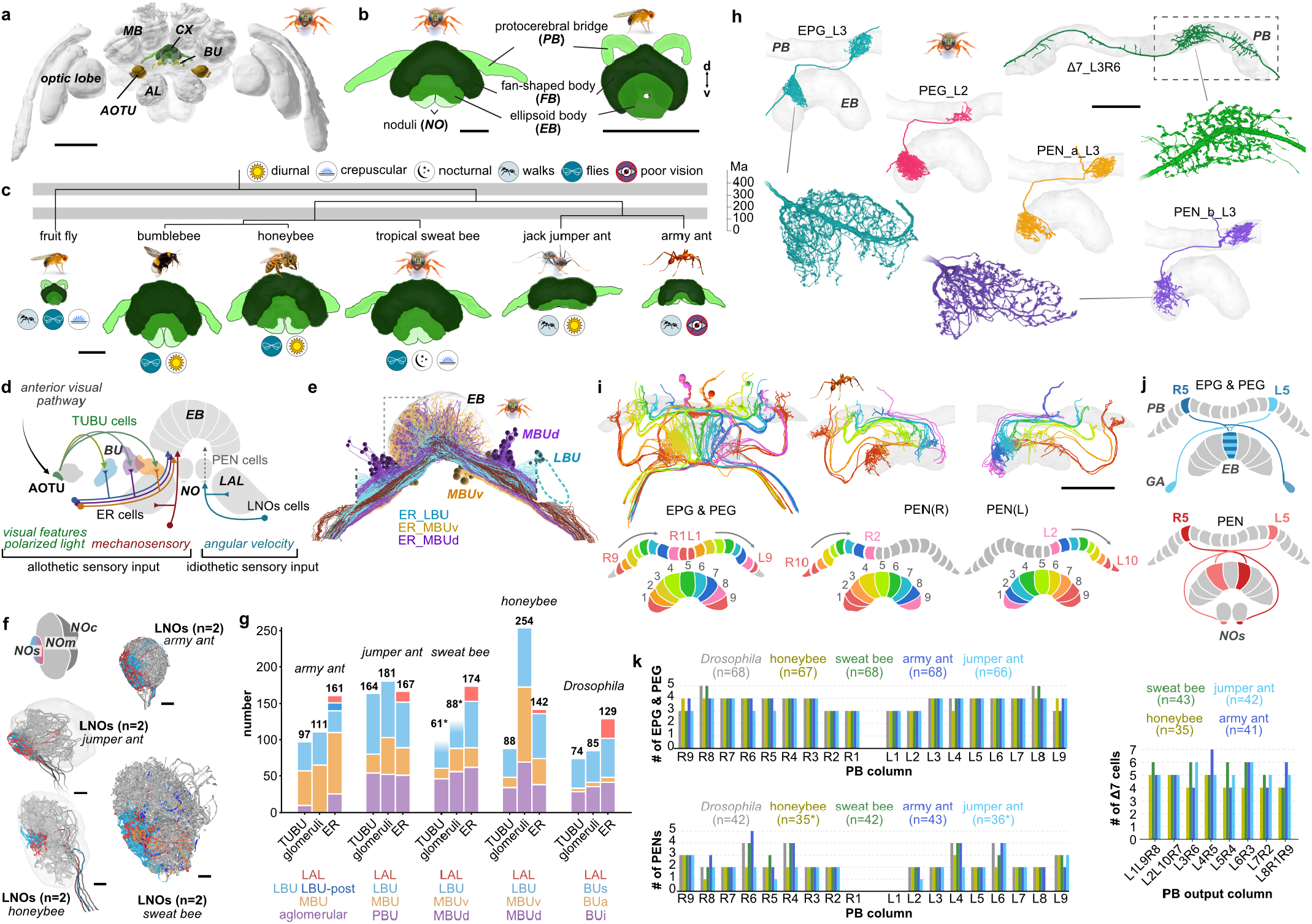
Repertoire of head direction neurons and their projection patterns in bees and ants. **(a)** The central complex in the context of the brain, illustrated by 3D reconstruction of a sweat bee (*Megalopta genalis*) brain. **(b)** The central complex in hymenopterans versus flies; shown are sweat bee (left) and *Drosophila* (right). **(c)** The central complex of all species examined: *M. genalis, A. mellifera, E. hamatum, M. nigrocincta*, complemented by the *Drosophila* and bumblebee central complexes. Icons illustrate key ecological traits. **(d)** Input pathways to the central complex, showing converging external (allothetic) and self-generated (idiothetic) cues. Ring neurons of the ellipsoid body (ER cells) relay allothetic visual and mechanosensory inputs from upstream regions (Anterior optic tubercle (AOTU), bulb (BU), lateral accessory lobes (LAL)) to head direction neurons in the ellipsoid body (EB). Lateral neurons of the noduli (LNO neurons) convey idiothetic rotational velocity signals from the LAL to PEN neurons in the noduli (NO). **(e)** All ring neurons of the ellipsoid body (ER cells) of the sweat bee. The inset highlights the synaptic-resolution image volume. Dotted vertical lines: border of image volume. **(f)** Organization of bee and ant noduli. Two LNOs neurons (colored) target the small unit of the noduli (NOs), LNOm neurons (gray) innervate the main unit (NOm), while the cap unit (NOc) was devoid of LNO cells in all four species. **(g)** Composition of the allothetic input pathways across species. Shown are numbers of AOTU projection neurons (TUBU), ER neurons of the EB, and bulb microglomeruli housing the giant synapses between both cell types. Colors: proposed homologies of parallel pathways across species. Asterisk: Lateral bulb inputs and glomeruli were partially outside the image volume. **(h)** Individual examples of the main cells comprising the core head direction circuit in bees: EPG, PEG, PEN_a, PEN_b and Δ7 neurons. Enlargements: details of branching trees. **(i)** Projection patterns of EPG/PEG and PEN neurons in the army ant. **(j)** Schematic of protocerebral bridge-ellipsoid body projection offsets for EPG/PEG and PEN neurons. **(k)** Quantities and distributions of EPG/PEG, PEN, and Δ7 neurons in all species. Scale bars: (a) 500 µm; (b,h,i) 100 µm; (f) 10 µm.

The central complex, as a brain region, is highly conserved^3, 4^ (Figure 1b). Like in flies, the central complex of other insects contains an array of repeating columns, formed by systematic wiring of different types of columnar neurons. Neurons homologous to all components of the fly head direction network have been identified across many distantly related insect species^32–37^, suggesting conserved overall circuit architectures. However, as the specific demands on such a circuit depend on the ecology of a species, we hypothesize that a species’ typical movement patterns, sensory environment, and navigational strategy provide evolutionary pressure to modify the functional properties of head direction coding. At the same time, strong pressure exists towards stability, ensuring continuous function of the head direction system. How evolution balances desired flexibility against required stability at the level of neural circuits is unknown.

To investigate how these seemingly conflicting demands are resolved, we have carried out volume electron microscopy based reconstructions of the head direction circuits in the central complex of four species of bees and ants. These insects are evolutionarily separated from flies by over 300 million years and exhibit diverse, advanced navigation behaviors with specific requirements on head direction coding (Figure 1c).

Our results revealed that the core set of cell types comprising head direction networks is conserved over at least 300 million years of evolution. While we found cell type identities and projection patterns to be highly similar across species, connectivity analysis showed that the feedback loops underlying the fly head direction circuit are fundamentally different in bees. However, computational models directly derived from connectome data demonstrated that both circuits effectively function as ring attractors. These circuits thus represent different stable circuit solutions to a shared computational problem - ensuring functional stability without the need to rigidly fix all circuit details. Flexibility is further facilitated by variability in the sensory input pathways upstream of the core head direction circuit, enabling functional specializations to evolve in line with ecological demands, without the need to modify key aspects of the computations required for head direction coding.

## The central complex of hymenopterans with different navigational strategies

We analyzed the head direction circuits of four hymenopteran species, all of which are established models for navigation research. Each species exhibits a distinct combination of navigational behaviors and sensory ecologies, allowing us to ask whether head direction circuits are adapted to different demands. These included a tropical sweat bee (*Megalopta genalis*), the honeybee (*Apis mellifera*), an army ant (*Eciton hamatum*), and the Australian jumper ant (*Myrmecia nigrocincta*; Figure 1c, Extended Data Figure 1a).

To reconstruct head direction circuits, we applied a novel data acquisition and analysis strategy^38^, in which we imaged an overview of the entire central complex of each species at cellular resolution and selected sub-volumes, from the same samples, at synaptic resolution (Figure 1c; Extended Data Figure 1f). For the synaptic-resolution image data, we exploited the repeating columnar layout of the central complex by placing imaging tiles following the typical trajectory of a columnar cell through the different central complex compartments (Extended Data Figure 1f). While we obtained high-resolution image volumes for all four species, we focused our dense neuron segmentation and synaptic-level connectomic analysis on the sweat bee and the army ant, the species with the highest quality image data. Reconstructing the four species using serial section block-face electron microscopy required imaging a total of 20,976 sections at 50 nm thickness, nearly three times the number used to image the entire nervous system of the fruit fly^39^.

To provide context for neuronal data, we first segmented the central-complex neuropils using our electron-microscopic image volumes. While the central-complex morphology was highly similar across species (Extended Data Figure 1d), its scale differed considerably, with the largest central complex in the sweat bee being roughly six times the volume of the smallest one in the army ant (Extended Data Figure 1e).

As a second step, we used the overview image data from all species to obtain cell numbers and general projectivity, focusing on neurons homologous to those that form the fly ring attractor network (EPG, PEG, PEN, and Δ7 cells; Figure 1h-k) as well as on neurons forming the sensory input pathways (ER cells; Figure 1d-g). To assign cell type identity, we used their arborization domains in the central complex: Neurons linking individual columns of the protocerebral bridge with the ellipsoid body were classified as EPG/PEG neurons, while neurons with an additional branch in the noduli were identified as PEN neurons. Within the protocerebral bridge, multi-glomerular neurons with dendritic branches in multiple columns and output in two to three individual columns were classified as Δ7 cells. After identifying all major cell types across all four species, we used the densely annotated, high-resolution data of *Megalopta* to confirm that PEG cells and both subtypes of PEN neurons (PEN_a, PEN_b) are also found in bees. Thus the full repertoire of cell types known from the *Drosophila* head direction circuit also exists in hymenopteran insects.

### Input pathways are highly evolvable

Which sensory cues are available and how informative they are for extracting directional information depends on the habitat an animal species occupies. While it is known that different insect species use different sensory cues for determining heading, it is not known how these preferences are reflected in the brain. We therefore first analyzed the sensory input pathways to the central complex across our species, asking to what extent these inputs are conserved.

In *Drosophila*, idiothetic (self-generated) and allothetic (external) signals reach the head-direction circuit via parallel pathways, terminating in the noduli and the ellipsoid body, respectively (Figure 1d). Idiothetic self-motion signals are conveyed by lateral neurons of the noduli (LNO cells), some of which are dedicated to carrying rotational self-motion signals (called GLNO cells in *Drosophila*). They drive columnar PEN cells to shift the EPG activity bump laterally^30, 31^, thus integrating angular velocity into an updated position of the head direction activity peak.

To avoid accumulation of angular errors, the position of the activity bump is also tethered to external sensory cues. These allothetic signals are provided by ring neurons of the ellipsoid body (ER neurons), which carry directional information about the visual panorama, polarized light, and mechanosensory information along multiple parallel pathways converging on the EPG cells^1, 40–44^. These inputs reach the ellipsoid body via the anterior optic tubercle (AOTU) and the bulbs. A key feature of this pathway is a set of giant synapses (microglomeruli)^40–43^ that are formed by AOTU projection-neuron axons (TUBU cells) and the dendrites of the ER cells. Individual microglomeruli serve as retinotopically arranged analyzer channels for low-level visual features^40^. The output of these feature detectors is provided to the EPG dendrites across all ellipsoid body columns via the wide-field axons of ER cells.

The overall layout of inputs described for the fly was conserved across all four hymenopteran species. Terminating in the noduli of the central complex, LNO neurons formed two parallel input channels. Their axonal branches delineated three noduli compartments: the small unit (NOs), the main unit (NOm), and the cap region, which was defined by the lack of LNO branches (Figure 1f). This tripartite organization was identical to that previously found in the bumblebee^37^. As no region devoid of LNO cells (cap region) exists in the fruit fly, the hymenopteran noduli showed a neuropil organization consistently distinct from that of flies. Despite this difference, the convergence of LNO input onto PEN columnar cells in the NOs was universally conserved, suggesting that the pathway encoding angular velocity is fundamentally required to integrate rotational movements for head direction coding.

While conserved in principle, the LNO/PEN system in the army ant stood out as distinct. In this species, the terminals of PEN and LNOs cells occupied a much larger relative volume of the nodulus (Figure 1f) and fiber diameters of PEN cells were dramatically increased (Figure 2a). Both findings suggest that the army ant head direction circuit relies relatively more on rotational self-motion signals, possibly compensating for less reliable allothetic visual inputs in these nearly blind insects.

**Figure 2:**
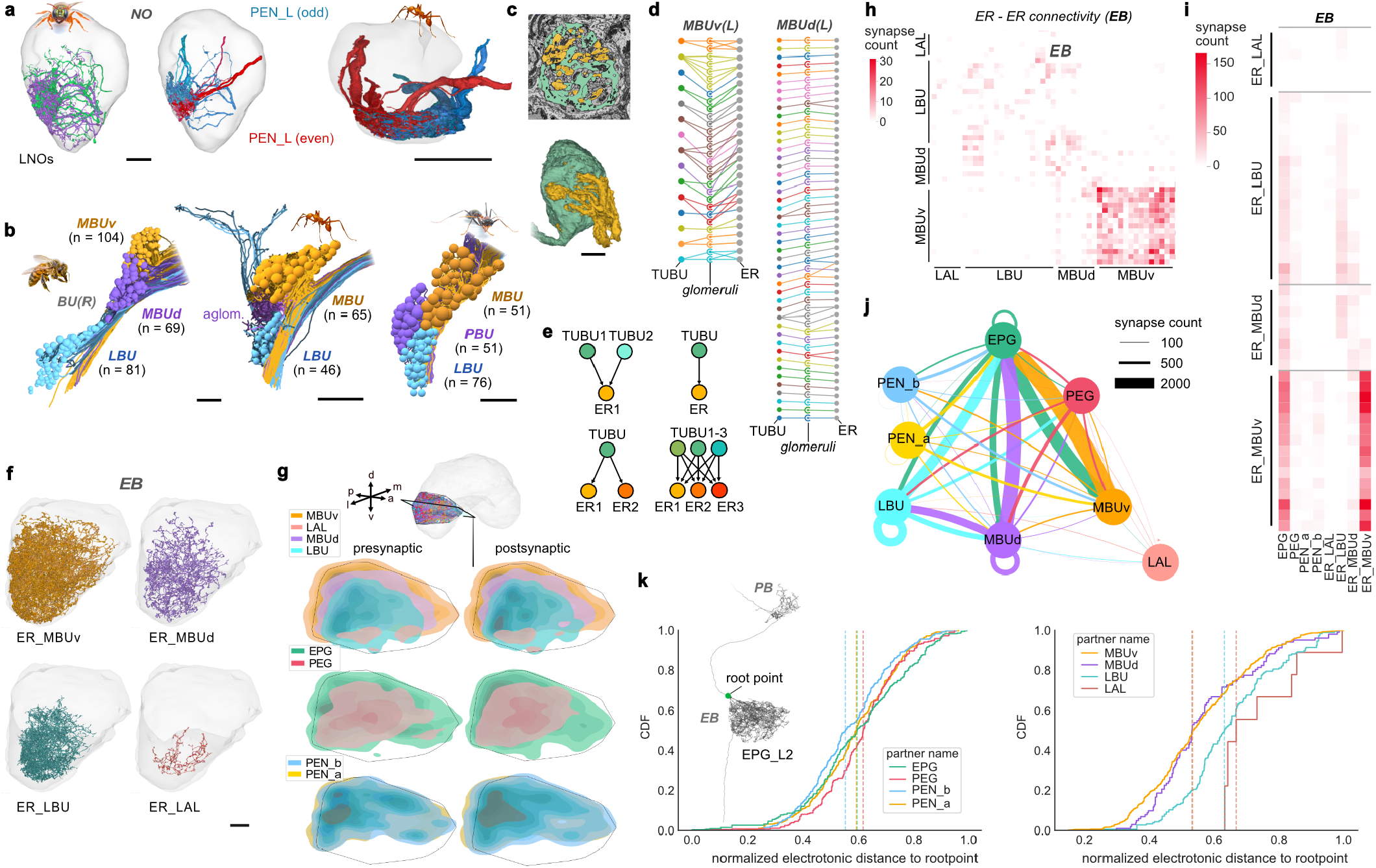
Convergence of sensory input pathways onto the head direction network across species. **(a)** Left: LNOs and PEN neurons in the sweat bee noduli. Right: PEN branches in the noduli of the army ant. **(b)** Organization of microglomeruli in the bulbs of the honeybee, army ant and jumper ant. Shown are microglomeruli (numbers indicated) and the postsynaptic ER cells. Note the unique, extended branching of some ER-LBU cells in the army ant. **(c-e)** Connectivity between TUBU and ER neurons in the bulbs. **(c)** Section and 3D rendering of one segmented microglomerulus of the MBUd region of the sweat bee. **(d)** Connectivity patterns between all TUBU and ER cells in the medial bulbs of the sweat bee. **(e)** Main connectivity motifs between TUBU and ER cells across species, showing different degrees of divergence and convergence. **(f)** Morphological reconstructions of ER neurons in the ellipsoid body, lateral view. **(g)** Distribution of pre- and postsynaptic terminals of head direction neurons innervating ellipsoid body column 2 in the sweat bee. d, dorsal; v, ventral; p, posterior; a, anterior; m, medial; l, lateral. **(h)** Adjacency matrix showing connectivity between all analyzed ER neurons of the sweat bee, illustrating the formation of intra- and cross-type networks between ER cells. Presynaptic neurons are on the y-axis. **(i)** Adjacency matrix showing connectivity between individual ER neurons and ER, EPG, PEG, and PEN neurons grouped by type in the sweat bee. Presynaptic neurons are on the y-axis. **(j)** Type-level connectivity graph of head direction neurons in the ellipsoid body (EB) of the sweat bee. To account for the only partial proofreading of ER cells, ER synapse counts for every ER partner were multiplied by the total number of ERs of that type. **(k)** Cumulative, normalized distribution plots of electrotonic distances between all input synapses and the putative spike initiation zone (green dot in neuron on the left) of ellipsoid body column 2 EPG neurons. Data were averaged across EPG_L2 and EPG_R8 neurons. Scale bars: (a) 20 µm (b) 3 µm (c) 25 µm (d) 20 µm

To examine whether similar patterns of differential investment also exist within the allothetic visual pathway, we investigated the organization of the inputs to ER neurons in the bulbs. Despite substantial differences in spatial arrangement at the neuropil level, the bulbs of all species were consistently divided into three compartments. While this neuropil configuration essentially resembled that of *Drosophila*, the suggested homologies remain to be confirmed by functional studies.

In each species, the three bulb regions contained distinct populations of presynaptic TUBU neurons (Extended Data Figure 3), different postsynaptic ER output cell types (Figure 2c), and variations in the connectivity ratios between input and output cells (Figure 2b). Microglomerular counts varied from 97 in the army ant to 257 in the honeybee (Figure 2c). Estimating from the number of microglomeruli, the repertoire of feature channels passing through the bulbs is larger in hymenopterans than in *Drosophila* (ca. 85). Even the almost blind army ant worker brain possessed more microglomeruli than the fly.

To ask whether the organization of the bulb might reflect a species’ ecology in more intricate ways, we extracted the connectivity motifs of each bulb region across our species (Figure 2b; Extended Data Figure 3). Generally, connectivity motifs between TUBU and ER cells varied from strict 1:1 connections and stereotypical 1:2 connections, to loose many-to-many connections that combined divergent and convergent motifs. These motifs were formed by a variable number of TUBU cells diverging onto a larger, but comparably constant number of ER cells (Figure 1g). The degree of divergence and the maintenance of input channel segregation correlated with the visual abilities of a species. This was particularly pronounced between the army ant (nearly blind) and the jumper ant (highly visual). In the army ant, the number of input neurons was reduced by one third, with the almost complete reduction of the posterior bulb pathway (Figure 1g; Extended Data Figure 3d). The residual components of the army ant posterior bulb did not show glomerular organization but large, overlapping neural branches, suggesting pooling of signals rather than feature extraction. Additionally, the lateral bulb exhibited many-to-many connectivity, suggesting cross-channel interactions despite the presence of microglomeruli. Finally, circa one third of the lateral bulb ER neurons possessed unique, extensive side branches in posterior brain regions outside the bulbs (Extended Data Figure 3d). While the upstream neurons of these cells are unknown, their branching pattern suggests that novel, non-visual inputs are recruited to contribute to allothetic head direction coding in the army ant - in line with their sensory ecology.

### A hierarchy of sensory inputs onto head direction neurons in bees

Across all species, the different populations of ER cells originate in distinct bulb divisions and target the ellipsoid body of the central complex. Thus, information originating from each population of TUBU cells is passed collectively to the ellipsoid body, but is shaped by the distinct connectivity patterns in the bulb giant synapses (Figure 1d,e,g; Figure 2b-e). We next investigated how these inputs target the dendrites of the EPG neurons at the synaptic level. To obtain comprehensive connectivity data for synaptic-resolution image volumes, we applied machine-learning–based segmentation, combined with manual proofreading, and automated synapse detection (see Methods; Extended Data Figure 1). Within the volume covered by our synaptic-resolution data, we reconstructed all arbors of a randomly selected sample of 10–15 ER cells for each type, representing 15% of their total population (Figure 2f).

In *Megalopta*, all reconstructed ER neurons connected to all EPG neurons present in our image volume and additionally formed strong connections to each other. These intra-type clusters were particularly pronounced for the ER cells receiving input from the ventral medial bulb, while more cross talk between lateral and dorsal-medial bulb neurons existed (Figure 2h,j). Beyond these overall patterns, the different ER types formed a clear hierarchy: ER neurons with inputs in the ventral medial bulb (ER_MBUv) formed the strongest connections onto EPG cells, followed by ER_MBUd and ER_LBU. This ranking is equivalent to the arrangement of these cells in the fly, where connection weights are strongest for input from the BUa, followed by BUi and BUs (Extended Data Figure 4b). The input weights onto EPG cells corresponded with the morphological arrangement of synaptic terminals in defined ellipsoid body layers (Figure 2f,g), where more strongly connected cell types are found in more dorsal-posterior layers. To assess whether this spatial arrangement of synapses has the potential to differentially impact the firing probability of EPG cells, we computed the electrotonic distance from the ER cells’ presynaptic terminals to the putative EPG cell spike-initiation zone. While all columnar neurons clustered tightly in this metric, ER types differed markedly, indeed reflecting their anatomical order (Figure 2k). Although the sensory features carried by these cells are largely unknown in the sweat bee, ER-MBUv cells were shown to strongly respond to polarized light^34^. As these cells provide most synapses onto EPG cells and synapse most closely to the putative spike initiation zone, they likely tether the bee’s internal compass strongly to the skylight polarization pattern, which is known to be essential for hymenopteran navigation abilities.

### Evolutionary hotspots within a conserved neuroarchitecture

Above, we identified the sensory input pathways to the central complex as highly evolvable. We next asked whether any aspect of the core head direction network stood out as an evolutionary hotspot. The most obvious difference between the *Drosophila* central complex and its hymenopteran homologue is the unique structure of the Drosophilidae fly ellipsoid body. While in *Drosophila* it represents a toroid with radially arranged segments, it is an open structure comprising a linear series of columns in bees and all other insects investigated to date (Extended Data Figure 1d). Consequently, to represent full 360° rotations, the circular network topology underlying head direction coding cannot be locally closed within the ellipsoid body. Assuming an equivalent coding scheme for head direction as in flies, once activity reaches one edge of an open ellipsoid body, it needs to jump to its opposite edge, if the animal continues rotating in the same direction. This activity jump must be enabled by recurrent loops that connect the edges of the ellipsoid body with the inner and outermost columns of the protocerebral bridge.

Our cross-species comparisons indeed identified these edge connections as an evolutionary hotspot, with different species presenting different solutions to the problem of how a topologically closed ring attractor circuit could emerge from a discontinuous anatomical structure. These differences manifested in structural, projection level, and synaptic-level changes (Figure 3a-h). Besides the distinct shape of the ellipsoid body, the lateral ends of the protocerebral bridge also showed structural modifications, comprising a tenth column in ants and bees. This additional neuropil compartment was particularly large in the army ant and, in both species, housed synapses of Δ7 and PEN neurons - forming connections that do not exist in *Drosophila* (Figure 3a,b, Extended Data Figure 5a).

**Figure 3:**
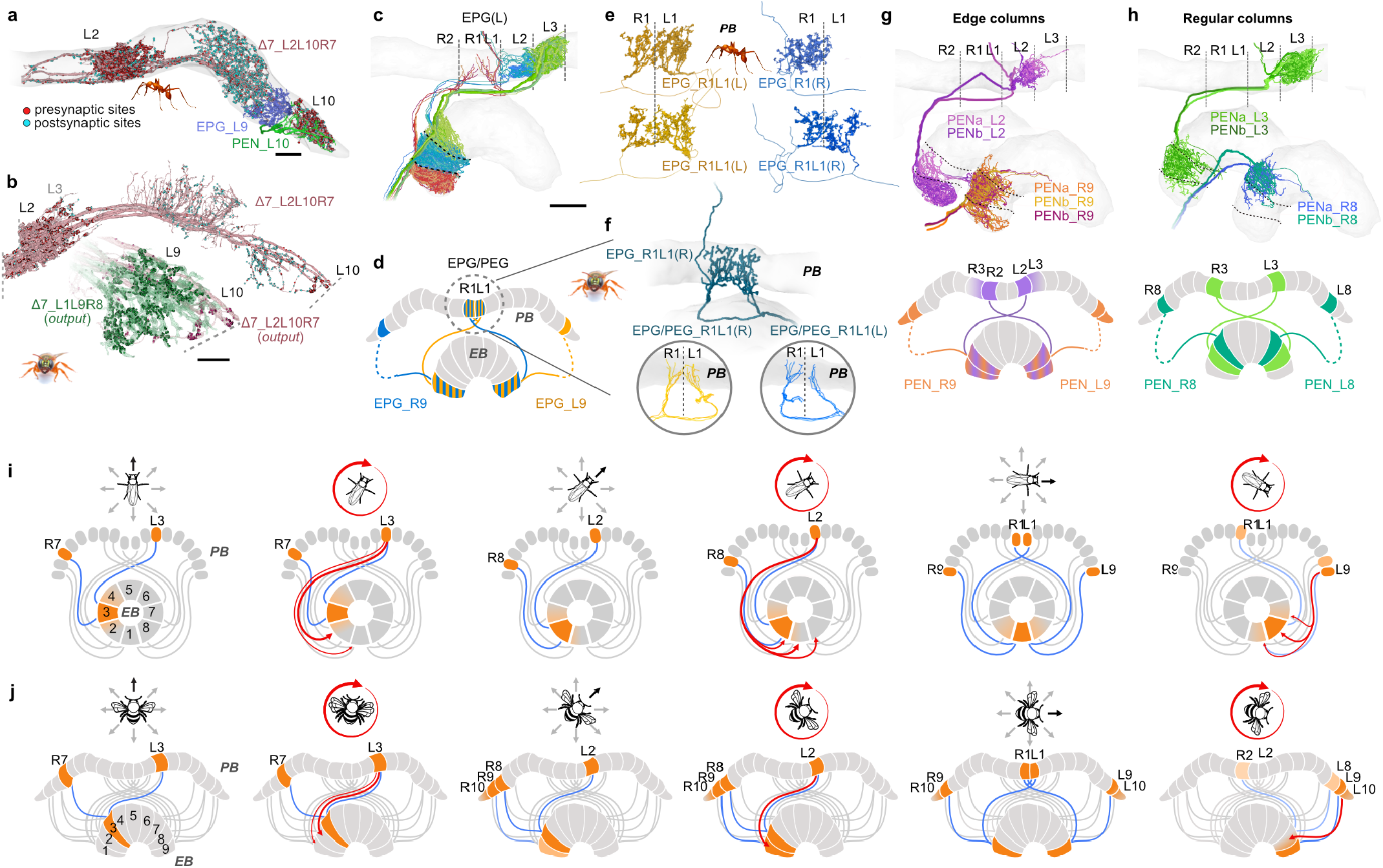
Highly evolvable features in the head direction circuit are suited to bridge a discontinuous ellipsoid body. **(a)** Δ7 neurons with the lateral-most output synapses form an additional (10th) column in the army ant (see Extended Data Figure 5a for fly homologue). Presynaptic terminals (outputs) are shown in red, postsynaptic terminals (inputs) in blue. **(b)** Sweat bee Δ7_L2L10R7 (red) and Δ7_L1L9R8 neurons (green) highlighting output synapses beyond the 9th column of the protocerebral bridge. Grey dotted line indicates image data boundary. **(c-f)** Projection domains of sweat bee and army ant EPG neurons. Ellipsoid body and protocerebral bridge columns are indicated by black dotted lines. **(c)** EPG neurons of three columns, including midline adjacent columns. **(d)** schematic representation of EPG neurons innervating the edges of the ellipsoid body. **(e)** Enlargements of midline projections of edge EPG neurons in the protocerebral bridge of the army ant. Note that all but one of these cells innervate both columns R1 and L1 evenly. **(f)** As (e), but for the sweat bee. The individual EPG neuron is a 3D reconstruction of a dye-filled cell (see Extended Data Figure 5b-d for fly and bumblebee neurons). **(g)** Projections of PEN neurons emerging from columns at the lateral edge of the ellipsoid body. Top, neural morphologies; bottom, schematic projections. **h** As (g), but for regular columns of the protocerebral bridge. **(i)** Illustration of how the EPG head direction activity bump is propagated across fused edges of the ellipsoid body during continuous turning in *Drosophila*. **(j)** Hypothesis of anatomically feasible mechanism by which the EPG activity bump could bridge the discontinuous lateral edges of the bee ellipsoid body during continuous turning. Scale bar: (a) 15 µm, (b) 10 µm, (c) 50 µm.

At the level of neural projections, differences were found in both the EPG and the PEN neurons. As in flies, PEN neurons in bees and ants do not innervate the midline columns of the protocerebral bridge (Figure 1k). To prevent a head direction activity bump from getting trapped in EPG neurons of this column, PEN neuron connections in the ellipsoid body need to close this gap. While in *Drosophila* this happens naturally because of directly overlapping projections at the fused ends of the fly ellipsoid body, in linear ellipsoid bodies, indirect measures have evolved. In *Megalopta*, PEN neurons from the outermost protocerebral bridge columns have twice as wide axonal fields in the ellipsoid body compared to all other columns, thus overlapping with EPG neurons two columns ahead instead of only one as for all other PEN cells (Figure 3g,h). This projection scheme thereby bridges the midline discontinuity of the PEN projections in the protocerebral bridge in a way distinct from *Drosophila* (Figure 3j).

All examined species, including flies, possess a set of EPG neurons that connect the midline-adjacent protocerebral bridge columns to the lateral edges of the ellipsoid body (Figure 3c-f; Extended Data Figure 5b). The details of how this is achieved differed across species, providing distinct modes and different degrees of cross-hemispheric coupling. Importantly, although EPG neuron projections were generally highly conserved across the studied insects, nearly each species showed a distinct configuration of midline EPG neurons, all of which differed from the fly (Extended Data Figure 5b). The multitude of circuit solutions indicated by our projectivity data suggests species-specific selection pressures, e.g. different demands on angular integration imposed by species-specific movement patterns. The higher rate of change at this position in the head direction circuit could even suggest the ring closure as a performance limiting feature, providing a possible access point to match circuit performance to species’ ecology - on top of adjusting sensory input pathways.

### A canonical head direction network

The sensory input pathways to the central complex target the network of columnar and local neurons of the ellipsoid body and protocerebral bridge, which, in *Drosophila*, generates a head direction signal. This signal is produced by computations that are largely determined by an anatomically defined network architecture formed by the stereotypical projection patterns of EPG, PEG, PEN, and Δ7 neurons. As the above analysis of evolutionary hotspots merely assumed that the involved neurons form a functional head direction circuit also in bees and ants, we next asked if this assumption can be grounded in circuit data.

Having identified all relevant cell types in our hymenopteran species, projectome-level reconstructions revealed highly conserved columnar projection patterns. Across our species, EPG/PEG neurons were organized in eighteen protocerebral bridge columns, converged on nine ellipsoid body columns (Figure 1i; Extended Data Figure 2), and followed a stereotypical projection pattern identical to that in the fly. PEN neurons were similar, but projected to an ellipsoid body column offset by one column from that targeted by EPG/PEG cells (Figure 1j). This offset is a defining feature of fly PEN neurons and enables the column-to-column shifts of peak neural activity during body rotations^30, 31^.

Cell numbers were nearly indistinguishable from those in flies, with generally four EPG/PEG neurons per column, two PEN cells, and five identical copies of Δ7 cells with identical columnar innervation patterns (Figure 1k; Extended Data Figure 2c,d). Interestingly, even the deviations from this general pattern were largely conserved, showing a smaller repertoire of columnar cells in the midline-adjacent columns and more PEN cells in columns four and six (Figure 1k). Moreover, the more highly resolved cell typing in *Megalopta* also confirmed that there were three EPG neurons complemented by one PEG neuron per column, and that the two PEN neurons found in each column belonged to different subtypes (PEN_a and PEN_b).

Together, these analyses demonstrate that the neurons comprising the fly ring attractor circuit are morphologically conserved across bees, ants, and flies, preserving key projection motifs over ∼350 million years of divergence^1, 37^.

Given the conserved morphology of head direction neurons across species, we assessed if the projectome-level conservation extends to synaptic connectivity patterns. Within our synaptic-resolution image volumes, we therefore reconstructed all branches of Δ7, EPG, PEG, PEN_a, and PEN_b neurons and characterized their connections across the *Megalopta* protocerebral bridge and ellipsoid body. This included three out of nine ellipsoid body columns as well as four out of nine protocerebral bridge columns in the contralateral hemisphere.

In flies, the connectivity between EPG, PEG, PEN_a, PEN_b, and Δ7 neurons yields a biological implementation of a ring attractor circuit^1, 2, 45^. At the core of this circuit are reciprocal, inhibitory connections between Δ7 cells, as well as two excitatory recurrent loops between the ellipsoid body and protocerebral bridge. The first loop enables activity to persist in absence of allothetic input (PEG-PEN-EPG loop) and the second one shifts the activity bump to a new position during angular movements (PEN-EPG loop).

In the bee, the general connectivity between cell types followed the canonical circuit identified in *Drosophila*. After receiving input from ER cells, EPG cells carry signals to the protocerebral bridge, where they converge onto Δ7 cells. These reciprocally interconnect with each other and provide output to the two other columnar cell types: PEG and PEN cells. Those project back to the ellipsoid body, where they connect to EPG cells, thus enabling two recurrent loops between the ellipsoid body and the protocerebral bridge.

Similar to *Drosophila*, these overall patterns are complemented by many local recurrences, seemingly redundant connections, providing additional routes for information and feedback connections. In combination, these significantly complicate the actual information flow within the fly and bee head direction circuits. Nevertheless, the overall connectivity motifs of cell-types in the sweat bee central complex matched those found in *Drosophila*, providing a conserved scaffold for generating, stabilizing and moving a head direction activity bump across the columnar array of the central complex (Extended Data Figure 3a).

### The inhibitory backbone of a biological ring attractor

To reveal the structure of the bee head direction circuit at the level of individual neurons, we first assessed whether the sweat bee Δ7 cells form the long-range inhibitory connections characteristic of the fly. Within the protocerebral bridge, we identified Δ7 neurons as the central hub of the network (Figure 4d). These cells receive input exclusively from EPG neurons and project broadly to all other columnar types, with their strongest outputs connecting back onto EPG neurons (Figure 4c,d). Additionally, they connect recurrently to other Δ7 neurons.

**Figure 4:**
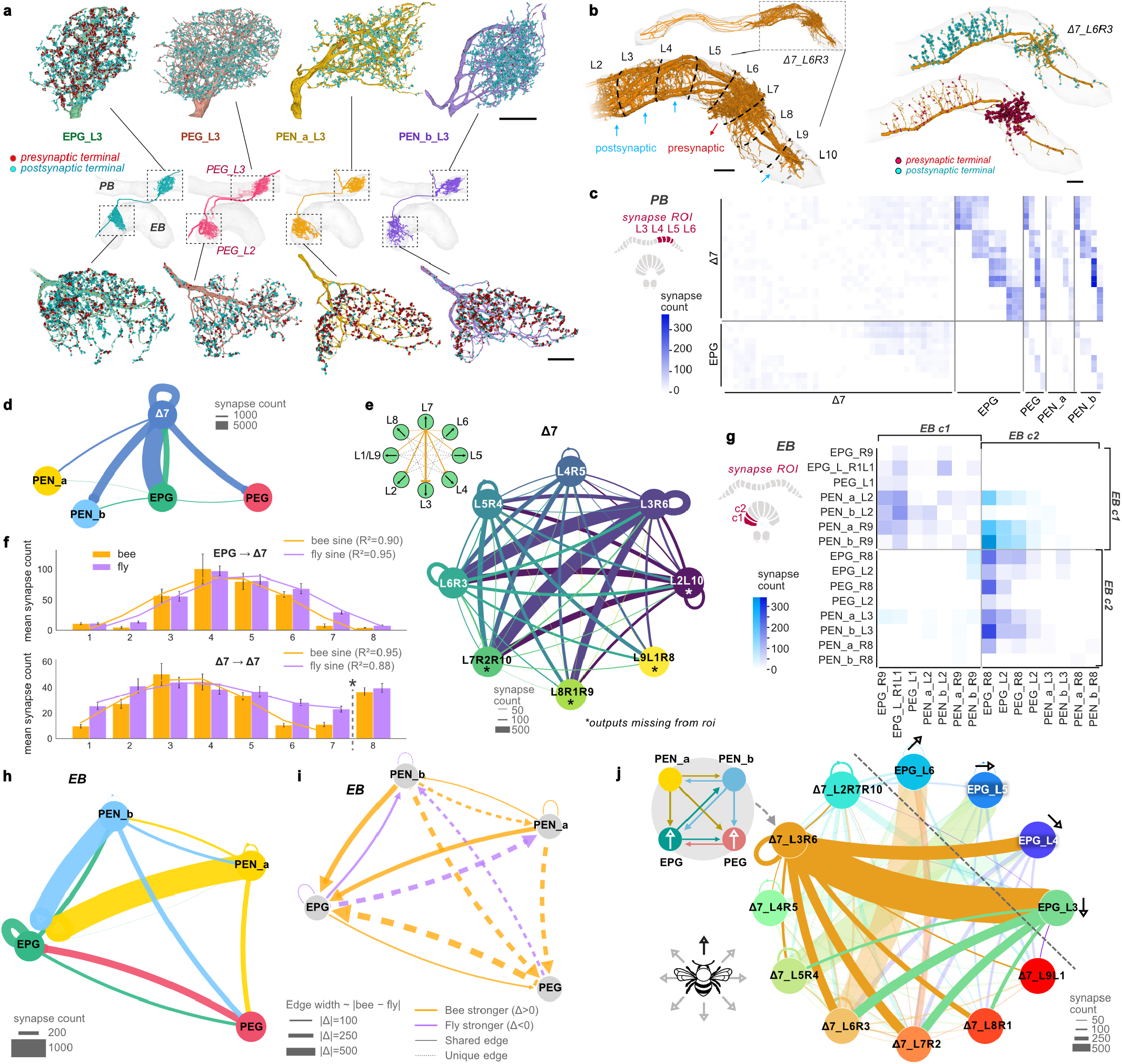
Ultrastructure of a biological ring attractor in the sweat bee central complex. **(a)** Representative morphologies of core head-direction neurons in the protocerebral bridge (PB) and ellipsoid body (EB), with pre- and postsynaptic sites. **(b)** Frontal view of example Δ7 neurons showing pre- and postsynaptic distributions. **(c)** Connectivity matrix of individual neurons in the PB; schematic: analyzed PB columns. **(d)** Cell-type connectivity graph for the PB (see also Extended Data Figure 6a-b). **(e)** Δ7 connectivity grouped by output column, revealing ring-attractor–like organization (idealized schematic at left). Asterisks: outputs outside the imaging volume. **(f)** Periodic organization of EPG and Δ7 connectivity across PB columns. Mean synapse counts (± SEM) for EPG → Δ7 (top) and Δ7 → Δ7 (bottom) connectivity in sweat bee (orange) and fly (blue); curves show best-fitting sinusoids (see Methods). Self connections (right of asterisk line) are not included in sinus fit. **(g)** Connectivity matrix of columnar neurons in the EB, grouped by EB column (c1 and c2). Light blue indicates cross columnar overlap of PEN projections. **(h)** Type-level connectivity graph of excitatory columnar neurons in the sweat bee ellipsoid body. **(i)** Difference graph highlighting species-specific connectivity among core excitatory neurons (see also Extended Data Figure 6c-d). An edge is defined as unique if it contains fewer than three synapses on average across all instances of this edge within our data volume. **(j)** Complete ring-attractor circuit. Top left, schematic of excitatory columnar connectivity (from **d**,**h**); main graph shows Δ7 and EPG neurons preferentially targeting PB columns representing directions 180°apart. Scale bars: (a-b) 20 µm.

As in *Drosophila*, synapses from EPG cells onto Δ7 neurons followed a sinusoidal distribution (Figure 4f), suited to reshape the EPG head direction signal into a sinusoid signal within the Δ7 neurons. The output branches of Δ7 cells are located four columns away from the center of their dendritic field (Figure 4e). This means that a head direction signal arriving on a Δ7 cell from a specific EPG neuron will cause inhibition of the EPG neuron with the opposite directional tuning (180° phase-shift) (Figure 4j). This pattern is suited to stabilize any current head direction activity bump. Similar to the EPG cell inputs, the within-type connections of Δ7 neurons also showed a sinusoidal profile across the population of Δ7 cells (Figure 4f). This pattern was more pronounced than the flatter distribution in *Drosophila*, mirroring the difference in dendrite density across the protocerebral bridge columns within any single Δ7 cell. Notably, the arrangement found in the bee is more similar to that reported by light-microscopical data from most other insects^18, 33–35, 46^.

Overall, if we assume that Δ7 neurons are also inhibitory in bees^18^, their synaptic connections are fully compatible with the idea that these cells form the systematic reciprocal inhibition required as the backbone of a ring attractor circuit for head direction coding. The difference in within-type connections of Δ7 neurons between flies and bees implies that the bee ring attractor, and possibly that of other insects, is maintained by a sinusoidally modulated reciprocal inhibition rather than a uniform, global inhibition supported by local excitation as in *Drosophila*, effectively implementing a different type of ring attractor^25^. This finding corroborates assumptions made by models based on projection level data that predicted distinct behavior of the head direction signal based on Δ7 neuron anatomy^34, 47^.

### Divergent core feedback loops at the heart of the head direction circuit

Unlike the largely conserved local circuitry in the protocerebral bridge, which forms the proposed inhibitory backbone of the ring attractor circuit, the feedback loops between the ellipsoid body and the protocerebral bridge were fundamentally different in the bee. This was surprising, as the involved cell types were indistinguishable between flies and bees at the level of projection patterns.

The first recurrent loop was formed by EPG cells and both types of PEN neurons. PEN neurons synapse strongly onto EPG cells in the ellipsoid body. Reciprocity is provided in the protocerebral bridge, where EPG cells form direct connections onto both PEN types, and additionally provide strong indirect input to PEN cells via Δ7 cells. Due to the one column phase shift between the protocerebral bridge and the ellipsoid body for PEN cells, this recurrent connection is suited to shift activity laterally along the ring attractor. This shifting loop is not only qualitatively identical to its fly counterpart, but also shows largely matching connection weights. Exceptions to this are the much stronger inhibition between Δ7 cells and all downstream PEN cells, as well as weaker local feedback connections between EPG and PEN neurons in the ellipsoid body (Extended Data Figure 6c,d).

The second recurrent loop is thought to stabilize the head direction bump in the absence of sensory input. It is formed between EPG and PEG neurons innervating the same columns. In the bee, these connections are directly recurrent, with EPG cells connecting to PEG neurons in the protocerebral bridge, and PEG neurons synapsing strongly onto EPG neurons in the ellipsoid body. Surprisingly, the latter connection is missing in flies. In *Drosophila*, it is replaced by a connection from PEG neurons to PEN_b neurons, a connection that is absent in bees. The direct EPG-PEG reciprocity in the bee stabilizing loop is thus replaced by an indirect connection via PEN_b in the fly (Figure 4h; Extended Data Figure 6c,d). Additionally, the majority of connections in the ellipsoid body have different weights between the two species, with connections involving PEG neurons generally being unique to, or stronger in bees.

Taken together, the different pattern of inhibition between Δ7 cells in the protocerebral bridge, the different setup of the stabilizing loop, and different weights of feedback connections suggest that the head direction circuit in bees functions differently from the well described fly ring attractor.

### Distinct circuits in bees and flies are functionally equivalent

To test whether the different implementations of head direction circuitry predict differences in performance, we generated detailed, anatomically constrained models of the bee and fly circuits. To extrapolate the entire central-complex connectome of *Megalopta*, we combined the obtained local connectomes with our global projectome (Figure 5a). We established local arborization kernels learned for each cell type from local connectomes and used them to extrapolate the missing connectivity of the global projectome. To validate this approach, we applied the extrapolation algorithm to artificially constrained connectivity data from the *Drosophila* hemibrain dataset^1^. Here, the extrapolated connectivity matrix was almost completely identical to the full connectivity matrix directly obtained from the hemibrain (Extended Data Figure 7).

**Figure 5:**
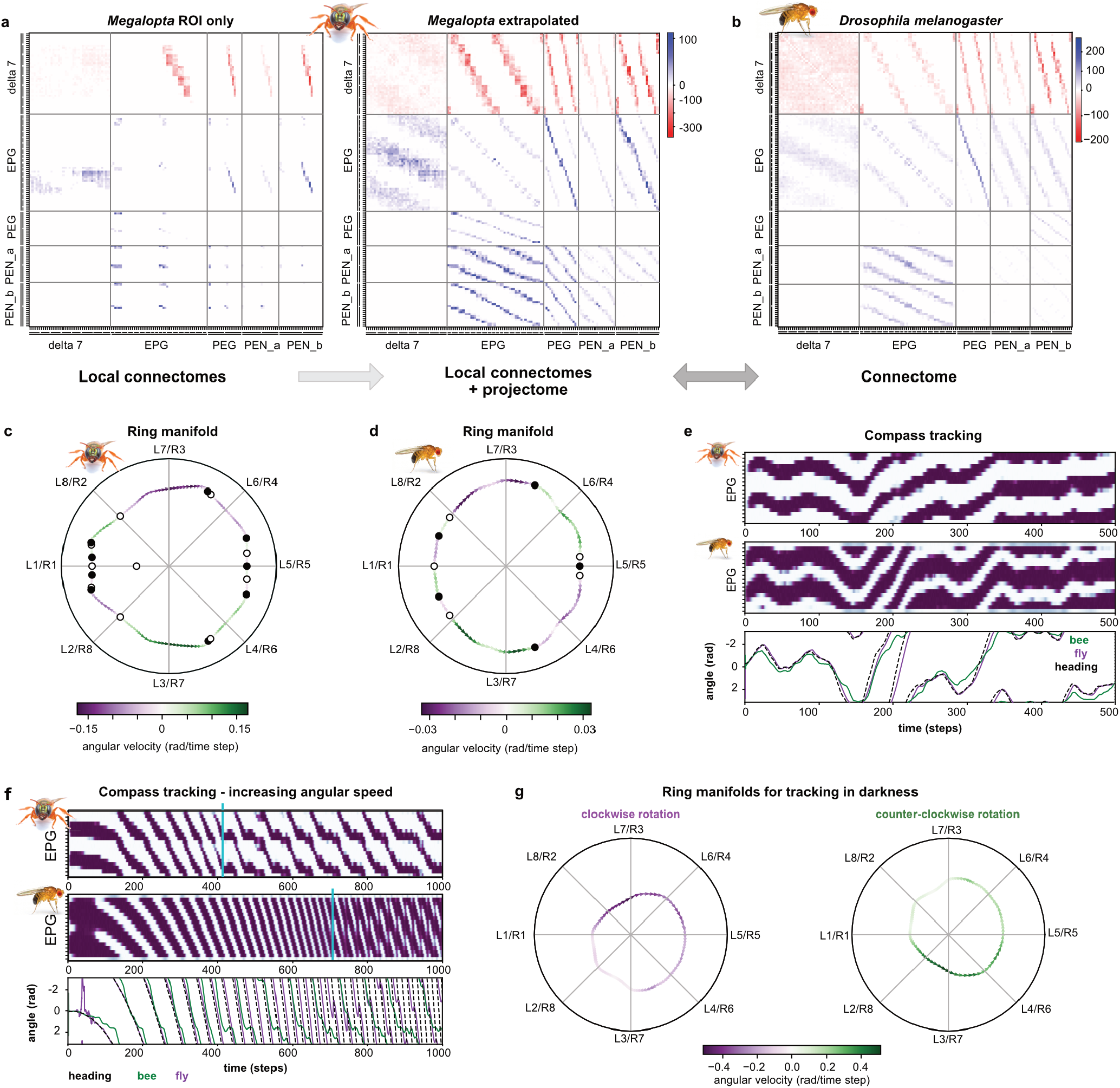
Comparative modeling of sweat bee and fly head direction circuit. **(a)** Adjacency matrix directly obtained from the synaptic-resolution image volumes of the *Megalopta* central complex (left). This partial connectome was used to extrapolate a full central complex connectome, using learned local arborization kernels combined with central-complex wide projectome data (right). **(b)** Adjacency matrix of the full *Drosophila* central complex, obtained from the hemibrain dataset. **(c**,**d)** Ring manifolds for neural activity in the extrapolated bee and fly circuits, demonstrating the existence of ring attractor dynamics resulting from the biologically constrained models in either species. Circular axis: anatomical space in central-complex columnar coordinates (population vector average projection). Shown are stable equilibria (filled circles), unstable equilibria (open circles), as well as local drift speeds of localized activity along the ring manifold. Slower drift speed indicates better approximation of an ideal ring attractor. **(e)** Localized activity bumps expressed in the EPG neuron population (shown for the protocerebral bridge) reliably track dynamically changing compass heading in the presence of visual input, both using extrapolated bee and fly models. **(f)** EPG-neuron compass tracking during systematically increasing rotational velocities show that the fly-based circuit enables following of faster rotational velocities compared to the bee based circuit. **(g)** Cyclic trajectories during imbalanced PEN neuron activation, mimicking rotational velocity input in the absence of visual input, shown for the bee based model. Inhibition of PEN neurons in one hemisphere results in smooth movement of the activity bump either in clockwise or counter-clockwise direction along the ring manifold.

In both systems, we simulated neural activity of EPG, PEG, PEN_a, PEN_b, and Δ7 neurons in a virtual agent performing rotational movements. Performance was quantified as the ability of the circuit to encode heading and keep in sync with imposed angular movements. These simulations were run either in darkness (only idiothetic input to PEN neurons) or with allothetic inputs as reference signals, directly relayed to EPG neurons.

All models were based on the synaptic weights derived from the connectome data in either species. To simplify and smoothen the biological data, multiple instances of identical neurons in any single column were grouped, and only the learned arborization kernels were used to implement the connectivity matrix. This enforced symmetry across the midline and eliminated local fluctuations of synaptic weights. The only other adjustments made were globally tuning inhibition versus excitation strengths, and uniformly adjusting background firing rate. To verify that our extrapolation algorithms capture all essential properties of the central complex connectivity, we performed the same simulations based on arborization kernels learned from the full ground-truth connectome of *Drosophila*, with identical results (Extended Data Figure 9).

Importantly, our simulations showed that both species’ circuits performed as ring attractors, encoding a single peak of neural activity that tracked angular movements (Figure 5c-e; Extended Data Figure 8). In both species, this was highly accurate when allothetic information was available. When testing the limits of how fast compass input can be tracked, the fly-based circuit was able to follow faster angular velocities (Figure 5f). Both circuits also tracked heading in darkness, using solely self-motion input (Figure 5g; Extended Data Figure 7). As in biological measurements^19^, angular errors accumulated in darkness, an effect that was more severe in the bee circuit compared to the fly circuit. Notably, across all situations both models encoded full, continuous 360° movements. Thus, in the bee model, the bump traversed from one lateral end of the central complex to the other, indicating that the bee head direction circuit is indeed functionally closed, despite possessing a morphologically open ellipsoid body (Figure 1b).

The emergence of ring attractors directly from connectomics data is remarkable, as the underlying feedback loops driving both circuits were fundamentally different (Figure 4i; Extended Data Figure 6). We conclude that both circuits represent different stable solutions to constructing a ring attractor circuit from homologous neurons. The existence of at least two viable circuit solutions relaxes the evolutionary pressure on stability, as it shows that within the complex parameter space provided by the multitude of neurons and synaptic connections of the central complex, reliable ring attractor dynamics can emerge in multiple ways. If these different solutions yield slightly different properties for direction encoding, the flexibility needed to match specific behavioral or ecological demands could readily emerge.

As the fly circuit model performance does not fully match the performance of the biological fly central complex as measured by calcium dynamics (e.g. in^19^), quantitative comparisons between the bee and fly models provide limited insights until more details of the biological circuits become known and are added to the model. To nevertheless illuminate the significance of the distinct connection patterns identified across both species, we next probed the susceptibility of both circuits to virtually ablating key components. We ablated Δ7, PEG, PEN_a, and PEN_b neurons and all combinations of these in both models and checked for the existence of ring attractor dynamics, the ability to track rotational movements in the presence of compass input, and the ability to shift the activity bump in response to simulated rotational self-motion (Extended Data Figure 10.

Four conclusions were drawn from those experiments (Extended Data Figure 10). First, without Δ7 neurons, no head direction bump exists across the central complex. Activity evenly spreads across the entire structure, highlighting the key importance of the conserved reciprocal inhibition provided by these cells to constrain and shape the head direction activity bump. Second, leaving only EPG neurons intact enables tracking of a simple visual environment, despite a complete lack of ring attractor dynamics and an inability to respond to self-motion cues, suggesting a possible ancestral starting point in the evolution of head direction coding. Third, both types of PEN neurons are required for ring attractor dynamics and for enabling angular integration of self-motion inputs. How much the overall circuit suffers from the loss of either PEN subtype was species-specific, suggesting that the different circuit layout across species changes the importance of the detailed feedback loops built by these cell types. Finally, ablating PEG neurons was least detrimental to both circuits, yielding intact ring attractors and an ability to follow self-motion inputs. However, the gain of self-motion responses, as well as the drift speed of the activity bump along the ring manifold, were increased to different degrees in either species. This implies that the temporal dynamics of the activity bump, i.e. the ability to remain stable or to respond to angular velocity inputs, strongly depends on the specific connections made by PEG cells. This finding strongly supports our anatomical data, where the connections made by these cells were found to be the most evolvable in the core head direction circuit. This indicates that PEG neuron connections, and to a lesser degree those of the PEN subtypes, can be adjusted during evolution to tune the dynamic properties of the ring attractor without breaking its fundamental properties.

## Discussion

Using volume electron microscopy, we have mapped the circuits homologous to the fruit fly head direction circuit across four species of bees and ants. The resulting cell type repertoire, projection patterns, and connectivity maps were then compared to the fly. We found that the architecture of this circuit is fundamentally conserved across all species in this study and must therefore have evolved approximately 350 million years ago^48^. This suggests strong selection pressure to maintain not only cell type identity, but cell numbers and detailed projection patterns, indicating functional conservation. This is in line with the fundamental need for head direction coding in any animal with directed behavior and suggests that once a functional circuit appeared early in insect evolution, it was maintained without major modifications. At least across holometabolous insects, an EPG-neuron centered compass circuit is most likely part of an ancestral navigational decision network, dating back at least to the end of the Devonian period. Nevertheless, each species possesses a distinct behavioral repertoire, sensory environment, and biophysically constrained body movements, creating specific demands on head direction coding. Consequently, any head direction circuit needs to adapt, for instance, to different sensory information available in its environment. How core brain circuits retain the capacity for evolutionary change, while remaining functional at all times, is an unanswered question across animals.

Within the insect head direction circuit we have identified several highly evolvable features, indicating that these complex circuits do not change uniformly, but contain hotspots for evolutionary change^49^. This suggests that small differences in key locations have performance-defining consequences without major detrimental effects on overall function. One such evolutionary hotspot was located at the sensory input side, where we identified anatomical evidence of input rerouting, different pre-processing principles, and addition of processing capacity. In conclusion, changing inputs of a highly conserved circuit appears to be an effective solution to compensate for changing environmental niches occupied by different species, as this strategy reduces the pressure for change on the core networks.

Yet, the dynamic properties of the core head direction circuit also need to be adaptable, considering the vastly different rotational speeds produced, for instance, by the ultra-fast flight saccades of a fly, compared to the much slower movements of a nocturnal bee or a walking ant. All of these movements have to be reliably captured by an internal head direction representation. To optimize this task, it appears highly advantageous to match the dynamic properties of the circuit with the behavioral abilities of a species. Hotspots of change identifiable at the level of projection patterns were sparse in our data. However, cell types that were otherwise highly conserved differed markedly between bees and flies at the level of synaptic connectivity. These changes were concentrated in cell types that form the core recurrent feedback loop of the attractor circuit, as well as the detailed layout of its inhibitory backbone^1, 2, 47^. Numerous unique connections, altered connection weights, and different synapse distributions across cell populations generate a distinct implementation of a ring attractor in the bee compared to *Drosophila*^25^. Nevertheless, both circuits constituted functional ring attractors when simulated in computational models that were directly derived from connectomics data. This finding is important, as it highlights that similar circuit function can emerge in at least two different ways from a complex set of homologous neurons, indicating some degree of degeneracy in neural circuit design. Our modeling data also showed that the core recurrent feedback loops contain elements that are not essential for maintaining overall functionality, but their virtual removal affects the dynamic properties of the ring attractor of the head direction network. This segregation between core features and tunable elements, enabled by the observed complexity in circuit layout, generates the flexibility that ultimately allows functional convergence. We predict that functionally convergent versions of head direction circuits have different dynamic properties and can hence drift into optima aligned with ecological demands of a species.

In general, connectomics has revealed neural architectures that appear much more complex than what had been expected based on functional data and earlier computational models^34, 50, 51^. Our results suggest that it is this complexity and the associated partial degeneracy that allows core circuits to evolve without breaking. The enormous complexity found in neural systems offers the chance to achieve similar computations in multiple ways, even without major changes in cellular composition or large-scale alterations of brain architecture. It might thus be evolutionarily advantageous to maintain seemingly redundant connectivity despite the energetic cost of sustaining it. While the evidence for this hypothesis is based on the insect head direction system, it is worth exploring whether similar principles could balance stability and flexibility in brains of other animals, including vertebrates.

## Methods

### Nomenclature

We adopted the *Drosophila* nomenclature for central complex regions and the corresponding abbreviated terms for cell types^1, 52, 53^. Although earlier studies in non-dipteran insects used an alternative naming convention, strong evidence for central complex homology across insects, together with the extensive functional and anatomical data available in *Drosophila*, makes this framework the most accessible for comparing our data to existing fly connectomics resources.

Given the stereotyped nature of the central complex, the projection patterns obtained from manual skeleton tracing were often all that was required to identify most cell types. Without connectivity data however, we were unable to distinguish between subtypes that innervate the same regions but differ in their polarity, such as is the case for EPG vs. PEG, for instance. We therefore grouped EPG/PEG neurons as EPGs, and PEN_a and PEN_b as PENs for all species for our projectome analysis.

In the sweat bee, dense reconstruction identified the columnar subtypes EPG, PEG, PEN_a and PEN_b. EPGs were distinguishable from PEGs primarily by polarity and, to a lesser degree, by morphology: EPGs possessed predominantly presynaptic terminals in the protocerebral bridge, whereas PEGs showed the opposite pattern (Extended Data Figure 4). However, all arborizations were mixed, i.e. contained both input and output synapses, and no clear polarity preference was observed in the ellipsoid body. PEN_a and PEN_b were separable by morphology alone, with PEN_a projecting across adjacent columns and PEN_b remaining column restricted (Figure 1h), consistent with the corresponding PEN subtypes in the fly.

### Animals

Neurons were reconstructed from the central complex of five species of hymenopterans: the honeybee (*Apis mellifera*), sweat bee (*Megalopta genalis*), army ant (*Eciton hamatum*), desert ant (*Cataglyphis fortis*, and jack jumper ant (*Myrmecia nigrocincta*; Figure 1I). Only foraging workers were used in this study.

Sweat bees and army ants were collected in the rainforest near the Smithsonian Tropical Research field station on Barro Colorado Island (Panama). *E. hamatum* workers were collected from foraging colonies and placed in ventilated vials containing cotton balls soaked with sucrose solution until being processed. To collect *M. genalis* sweat bees, a light trap was set up around twilight when they are most active. Light traps consisted of a large white sheet illuminated by a light source containing UV light. Captured sweat bees were kept in a vial containing cotton balls soaked in sucrose and water solutions until processing within a week of capture.

Honeybee workers were collected from a non-captive honeybee colony, maintained on the terrace of a building at the University of Queensland (Brisbane, Australia). Workers from this colony were allowed to forage freely on surrounding vegetation. Foraging *M. nigrocincta* jack jumper ants were collected from a nest located at the Sunshine Coast and transported to the University of Queensland where they were immediately processed for histology.

*C. fortis* ants were maintained in nests made of porous concrete (42 x 20 x 7.5 cm) that contained chambers of various sizes. The ants were allowed to forage in a square Plexiglas arena (56 x 56 x 10 cm). Lighting in the arena was provided by 120 cm long fluorescent tubes (36 W) with a distinct UV component. The day-night rhythm was set to 14-10 h from spring to autumn and 12 - 12 h in winter. All animals were exposed to natural skylight for at least three days to three weeks during behavioral experiments.

### Histology for SBEM

For SBEM histology, honeybees were stained using a standard protocol as described below. After acquisition of the honeybee dataset we found superior staining results using the method outlined in Hua et al. (2015)^54^, and therefore applied a slightly modified version of this protocol to the remaining species^38^.

For all five insect species, we began by anesthetizing the animal over ice for 3-5 min before removing the head directly into freshly made fixative. For the desert ant and honeybee, we used a standard EM fixative containing 2.5% glutaraldehyde and 4% paraformaldehyde in 0.1M sodium cacodylate buffer. At a later point, we discovered that using significantly lower fixative concentrations yielded insect brains with far better preservation. We therefore switched to using a fixative containing 0.75% glutaraldehyde and 0.75% paraformaldehyde in 0.1M sodium cacodylate buffer for the army ant, sweat bee, and jack jumper ant. For all species, the brain was immediately dissected away from the head capsule in fixative, removing the optic lobes, subesophageal ganglion, and the neural sheath whenever possible to aid with fixative permeation. All species were then fixed over night at 4°C. The following day, insect brains were rinsed 3 x 10 min in 0.1M sodium cacodylate buffer and then stored in the same buffer until further processing. While storage in buffer ranged from a few days to weeks for the honeybee and jack jumper ant, the remaining species were processed several months following fixation without any clear detriment to tissue quality.

At the time of processing, all insect brains except for the honeybee were placed in a solution containing 2% osmium tetroxide in 0.1M sodium cacodylate buffer and then microwave treated (Biowave, Ted Pella) using two 1 min on/1 min off/1 min on steps at 150 W under vac. Brains were then microwaved using 20 sec pulses with 20 sec rest periods at 250 W three times, swirling the brains in solution between pulsations. Brains were next left to incubate in the same solution for 55 min at room temperature before being transferred to a new solution containing 1.5% potassium ferricyanide, then microwave treated using the same pulse settings as described above and left at room temperature for 1 h. For the honeybee, fixed brain tissue was immersed in a solution containing 2% osmium tetroxide with 1.5% potassium ferricyanide in 0.1M sodium cacodylate buffer and subjected to the same microwave pulsation steps as described above. Honeybee brains were then left to incubate in solution for 45 min at room temperature.

All insect brains were next rinsed 4 x 5 min in ddH_2_O and then placed in a solution of 0.1% thiocarbohydrazide in ddH_2_O and left to incubate in the oven at 40°C for 45 min, omitting microwave pulsation steps. Following this, brains were transferred to fresh vials containing ddH_2_O and again rinsed 4 x 5 min with ddH_2_O. Tissue was then placed in 2% osmium tetroxide in ddH_2_O, subjected to a single microwave pulsation step at 1 min on / 1 min off / 1 min on (150 W) and left on the bench for 30 min. Samples were next rinsed 4 x 5 min in ddH_2_O and transferred to 1% uranyl acetate in ddH_2_O. All insects except for the honeybee were then left to incubate overnight at 4°C. Honeybee brains were left in uranyl acetate solution for 2 h and then rinsed 4 x 5 min in ddH_2_O before immersion in lead aspartate solution (described below).

The next day, non-honeybee brains still in uranyl acetate solution were transferred into an oven and heated to 50°C, where they were left to sit for 70 min. Samples were then removed from the oven, cooled to room temperature, and then rinsed 4 x 5 min in ddH_2_O. A solution of lead aspartate was made by adding 0.066 g lead nitrate to 10 ml of 0.03M aspartic acid in a falcon tube and pH adjusted to 5.5 with potassium hydroxide. The solution was heated at 60°C for an hour to aid dissolution. All insect brains were then soaked in lead aspartate solution for 45 min in the oven set to 50°C and then rinsed in ddH_2_O 4 x 5 min.

Lastly, brains were dehydrated through an ethanol series (20%, 50%, 70%, 90%, and 100%) at 15 min steps, transferred to acetone for 10 min, and then over the course of 2 hours embedded in Durcupan polymer (Sigma-Aldrich) at increasing ratios beginning from 3:1 (acetone:Durcupan), 1:1, 1:3, and finally twice into 100% Durcupan. Samples were then left to polymerize at 60°C for 48 hours. Polymerized samples were trimmed manually and mounted onto VolumeScope (Thermo Fisher Scientific) SBEM stubs using conductive epoxy. Brains were oriented so they could be sectioned at a horizontal orientation (Figure 1K) once in the SBEM.

### SBEM image acquisition

Volumetric EM images were acquired with a VolumeScope SBEM using MAPS 3 software (Thermo Fisher Scientific). During acquisition, all samples were imaged in low vacuum (10 Pa) using a VolumeScope directional back-scatter detector (VS-DBS; Thermo Fisher Scientific). Images were acquired using a landing beam energy of 2 kV, a beam current of 0.1-0.2 nA, and pixel dwell times of 2-3 µs. All samples were sectioned at 50 nm thickness, requiring a total of 20,976 sections to be cut and imaged for the combined four species (section counts: sweat bee, 6141; honeybee, 7222; army ant, 3870; jumper ant, 3743).

A multi-resolution approach was used which enabled imaging of the entire central complex of each species at cellular resolution (40-50 nm pixel scale) along with specific central complex compartments at synaptic resolution (8-11.5 nm; Figure 1J-K). The pixel scale in x and y was adjusted for each species to account for differences in brain size (see Extended Data Figure 1e). For the high-resolution images, pixel scale was set to ensure that synaptic features, such as vesicles and synaptic clefts, were clearly visible.

Central-complex compartments captured at high-resolution are schematized in Extended Data Figure 1f for each species. In the sweat bee and army ant, synaptic-resolution tiles were placed such that they covered nearly all of the columns in one hemisphere of the protocerebral bridge and 2-3 columns in the contralateral hemisphere of the fan-shaped body (FB), the ellipsoid body (EB), the right nodulus (NO_R), and, in the army ant and desert ant, the right bulb (BU_R; Figure 1J). This mapping follows the trajectory of a typical central complex columnar cell projecting from the left central complex hemisphere into the right.

### Multi-resolution image alignment and neuropil segmentation

To trace neurons from the overview (cellular resolution) dataset into the high-resolution datasets, all images must be aligned and registered to each other in 3D space. We began by aligning the overview image stack which served as a template for the high-resolution image stacks. All images were aligned using TrakEM2^55^, an open source plugin available with FIJI^56^ (ImageJ). To align the overview image stack, image tiles were first stitched together in x and y and then aligned slice by slice in z using SIFT^57^ (Scale-Invariant Feature Transform) linear translational transformations only. After alignment, overview images were filtered using a Gaussian blur with sigma 0.7 and then contrast adjusted using CLAHE.

To align the high-resolution image stacks, we extracted the relevant slices from the aligned overview image stack, cropped them, then imported them into TrakEM2. To adjust for differences in pixel scale, the overview image slices were scaled up to match the pixel scale of the high-resolution stack (i.e., when aligning an image with 10 nm pixel scale, a 40 nm overview would be scaled up by a factor of 4 in x and y). Next, the high-resolution images were affine aligned using SIFT to the overview stack, which served as a template. High-resolution images were then filtered and denoised in the same manner as the overview image stack. Images were exported as .tiffs for backup and as .jpeg image pyramids for reconstruction. 3D coordinates for each image stack were extracted by manually aligning a high-resolution image to an overview image using the Transform Editor in Amira (Thermo Fisher Scientific). Only a single image is needed during this process because each slice is 50 nm in Z. The Z coordinate can therefore be calculated by multiplying the slice number of the overview slice in which the high-resolution image stack begins by 50 nm.

To segment neuropils, overview image data was downsampled to 0.4 x 0.4 µm pixel scale in FIJI and then manually reconstructed using Amira’s segmentation editor. In the editor, the neuropil was filled in using the paint brush tool in 3-4 images spaced evenly apart from one end of the neuropil to the other. This was repeated in xy, yz, and xz, creating a scaffold around the neuropil. The Wrap Tool was then used to interpolate a smooth and accurate surface around the structure.

### Manual neuron reconstruction

Neuron skeletons were manually traced using CATMAID, a collaborative, open-source, and web-based tracing software^58^. Neurons could be traced seamlessly from the overview image stack into high-resolution image stacks thanks to a plugin incorporated from Hildebrand et al. (2017)^59^. Skeletons were traced by scrolling through image data while placing nodes that follow the trajectory of a cell’s fibres. Synapses were identified in the high-resolution image data by the combined presence of an active zone containing synaptic vesicles with either a synaptic cleft or postsynaptic specializations. We did not identify any structures that unambiguously corresponded to the T-bars reported in the fruit fly^53^. Gap junctions were not identifiable in our image data. Screenshots of neurons used in all figures were generated using the 3D Viewer widget in CATMAID.

Neuron skeletons and synapses were reviewed by expert tracers using the Review widget and following the methodology outlined in Schneider-Mizell et al. 2016^60^. To avoid redundant tracing, reviewers focused on the neuron fragments that contained the largest proportion of branches. Additionally, we identified 20 µm cubes in the high-resolution datasets that spanned areas of high variability in image quality for each neuropil and reviewed every individual fiber within these cubes. Lastly, we took advantage of the central complex’s bilateral symmetry to compare same type neuron morphology across hemispheres, as well as between homologous neurons across species, focusing our review efforts wherever divergent morphologies were identified.

Connectomic tracing of nervous tissue is time consuming and resource intensive. To make manual reconstruction of large volumes across multiple species feasible, we used a sparse, but targeted tracing approach. Using the overview image data, we traced the backbones and main fibers of cells to obtain quantities and general projectivities.

### Automatic image segmentation

We automatically segmented neurons within high-resolution datasets using a convolutional neural network (CNN). CNNs were trained to predict voxel-wise nearest-neighbor affinities and local shape descriptors using the method described in Sheridan et al. (2023)^61^. For ground-truth, we used the python library napari^62^ to densely reconstruct neurons and mitochondria from several 5 µm cubes of high-resolution data extracted from the *Megalopta genalis* dataset. Models were trained using gunpowder (https://github.com/funkelab/gunpowder/tree/main) and pytorch^63^ with the multitask LSDs (MTLSD) architecture described by Sheridan et al. (2023)^61^. We randomly selected hyperparameters for data augmentation and evaluated models by computing variation of information (VOI) against expert skeleton annotations, in 10 µm cubes not overlapping with training cubes. The best performing model was used for automatic segmentation of neurons for all high-resolution image data.

We used the CAVE engine^64^ to manually proofread segmentation. Volumes were artificially split at regular intervals to account for numerous merge artifacts caused by holes in neuron membranes at neuropil boundaries. This ensured that predictable merge errors were isolated, thus minimizing the number of split operations needed in favor of merge operations which are quicker to perform by users. For a more detailed account of our segmentation and reconstruction methods, including all code and machine-learning models, see^38^.

### Synapse detection

Automated prediction of synapses was performed using the ‘synful’ python package^65^. For ground truth, we annotated several 5 µm cubes from the image data volumes. These were used as training data for 16 models, each with the same architecture but differing image augmentation parameters. Performance of each was evaluated on a cube of image data that was not used in training, and outputs were compared to expert annotations on the same cube. The best performing model was selected and used to provide synaptic predictions for all image data. F-scores were calculated across several regions from each of the three datasets (PB, FB-EB, and NO) using the same method as used in^65^. For the PB, the average F-Score was 0.60, for the FB-EB it was 0.58, and for the NO it was 0.57. These scores are averaged across regions with varying image quality, but scores for best image quality were 0.63, 0.71, and 0.52 in the PB, FB-EB, and NO respectively. All F-scores are comparable to F-scores achieved in the *Drosophila* dataset with the same method. A threshold was applied to all synapse predictions to remove low-confidence predictions. F-score data is not limited to synapses where both cells are completed and identified, but the final data are.

### Analysis of connectomics data

All analysis of connectomic data was carried out in Python^66^ using custom scripts together with standard scientific computing libraries. Synapse tables were processed using pandas^67^ and NumPy^68^, with neuron–neuron connectivity represented as adjacency matrices derived from synapse counts between identified presynaptic and postsynaptic neuron pairs. Node connectivity graphs were generated from adjacency matrices using NetworkX. Graphs were treated as directed, weighted networks, with edge weights corresponding to synapse counts and colored by presynaptic identity. Only connections exceeding a minimum threshold of three predicted synapses between neuron pairs were retained. All visualizations of node graphs and adjacency matrices were generated using matplotlib^69^ and seaborn^70^.

For comparative analyses with *Drosophila*, we used the hemibrain connectome^53^, as central-complex neurons in this dataset have been extensively proofread and annotated, and it is currently the only connectome for which a comprehensive analysis of the central complex is available^1^. *Drosophila* hemibrain data were obtained from neuPrint^71^ using the neuprint-python package (https://github.com/connectome-neuprint/neuprint-python). To enable direct comparisons between the fruit fly and sweat bee, fruit fly hemibrain connectivity data were further filtered to match the ROIs available in the sweat bee dataset. Specifically, ellipsoid body and protocerebral bridge columns were defined using EPG neurons. EPG synapse coordinates were used to generate convex hulls with the SciPy^72^ and Trimesh (https://github.com/mikedh/trimesh) Python packages, and only synapses located within PB columns L3-L6 and EB columns c1-c2 were retained for analysis.

Neuron morphologies were visualized and analyzed using navis^73^, Neuroglancer^74^, CloudVolume^75^, and CATMAID^58^. Skeletonized neuron reconstructions were loaded into navis for quantitative analysis and visualization, while Neuroglancer and CloudVolume were used for interactive inspection of image volumes and segmentations. CATMAID was used for annotation, proofreading, and verification of neuronal morphologies.

To quantify periodic structure in population-level connectivity profiles, sinusoidal fitting was performed on column-wise summary statistics of Δ7 and EPG adjacency matrices (Figure 4f). For each profile, synapse counts were first summarized across observations to yield a mean connectivity value per anatomical column. These values were then modeled using a sinusoidal function:

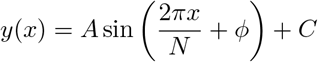

These data were then fit using non-linear least-squares optimization (*curve_fit* ^72^). Here, *A* denotes amplitude, *ϕ* phase offset, *C* a constant offset, and *N* the number of columns. Model goodness-of-fit was quantified using the coefficient of determination (*R*^2^), computed from residual and total sums of squares. Initial parameter estimates were derived from the empirical range and mean of the data.

To estimate the spatial organization of ER outputs, we calculated electrotonic distances from presynaptic sites to the putative spike initiation zone of each EPG neuron innervating ellipsoid body column 2 (EPG_R8 and EPG_L2; Figure 2k). Segmented neurons were skeletonized using the Python package pcg_skel (https://github.com/CAVEconnectome/pcg_skel.git). Fiber diameter estimates were derived from the neuronal mesh and retained as radii at individual skeleton nodes. The putative spike initiation zone was defined as the posterior-most branch point on the primary neurite as it leaves the posterior region of the ellipsoid body.

After identifying this reference point, electrotonic distances between each synapse and the spike initiation zone were computed using functions from the Navis package^73^. Because neurite diameter varies along the arbor, the length constant, *λ*, was calculated independently for each skeleton edge, with edge diameter taken from the radius of the terminal vertex. Each edge was then assigned an electrotonic length equal to its physical length divided by its corresponding length constant. The total electrotonic distance from a synapse to the spike initiation zone was obtained by summing these normalized edge lengths along the skeleton path connecting the two points. Thus, for a path consisting of edges with geodesic lengths *w*_1_, *w*_2_, …, *w*_*n*_ and length constants *λ*_1_, *λ*_2_, …, *λ*_*n*_, the electrotonic distance was calculated as 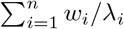 (see also^1^).

To generate synapse density plots (Figure 2g), ellipsoid body and ellipsoid body column 2 meshes were retrieved from CATMAID using pymaid. Synapse coordinates were obtained from the Megalopta central complex synapse table, restricted to synapses assigned to the ellipsoid body, and converted from voxel coordinates to CATMAID coordinates using the image-stack resolution. Presynaptic and postsynaptic sites located within ellipsoid body column 2 were identified using navis.in_volume. To standardize the projection of synapse positions, all synapse coordinates and mesh vertices were first centered on the ellipsoid body centroid. Principal component analysis was then fit to the combined ellipsoid body mesh, column 2 mesh, and synapse coordinates using scikit-learn. Coordinates were transformed into this PCA-aligned space, rotated by −15^°^ about the first principal component to obtain the desired viewing orientation, and projected onto the PC1-PC3 plane. The outline of column 2 was defined from the projected mesh vertices using a convex hull calculated with SciPy. Synapse distributions were then visualized as two-dimensional kernel density estimates using seaborn, with densities plotted separately for presynaptic and postsynaptic sites and grouped by neuron type.

### Model connectivity and assumptions

We compared two anatomically constrained models of the head direction circuit: one extrapolated from sweat bee connectivity and the other from the fruit fly hemibrain connectome (Figure 5a; Extended Data Figure 8).

### Simplifications

1. Neurons were grouped by column type (cell type and protocerebral bridge column innervation; see Nomenclature). The grouped connectivity was computed as the mean number of synapses between an individual pair of cells in each pre-synaptic and post-synaptic group. This implies assuming a compensation mechanism for varying numbers of cells per group.
2. The following fly Δ7 neurons were merged: Δ7_L4R5 & Δ7_L4R6, Δ7_L5R4 & Δ7_L6R4, and Δ7_L6R3 & Δ7_L7R3.

### Reconstruction of full sweat bee connectivity

To reconstruct the full sweat bee connectivity matrix, we assumed that the number of synapses between two cells is a function of the overlap of their arborizations. This was modeled as ‘arborization kernels’ describing how a cell of a specific type arborizes around the center of its projection (the latter can be determined from our low-resolution data).

Let *C* denote all cells in the (grouped) connectome to be reconstructed. The estimated number of synapses between cells *i, j* ∈ *C* in region *r* ∈ *R* = {EB, PB} is denoted 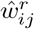. Additionally, we use the following notation:

- *t*(*i*) ∈ *{*Δ7, EPG, PEG, PEN_a, PEN_b*}*
- *L* is the set of columns in the EB and PB.
- *L*_*r*_ ⊂ *L* is the set of columns in ROI *r*.
- *P*_*r*_(*i*) ⊂ *L* is the set of projections of cell *i*.
- *δ*(*η, ν*) is the offset/signed distance between two columns *η* and *ν* in the same neuropil.

Given an output kernel 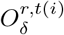 for a presynaptic cell *i* with column type *t*(*i*) in the ROI *r*, and an input kernel 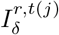 for postsynaptic cell *j*, we modeled the synapse count between cells *i* and *j* in *r* as

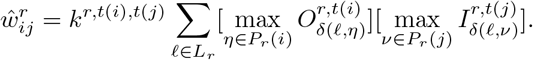

In other words, each distinct projection location adds to the total arborization of the cell, and the sum of the column-wise product of the output and input arborizations is proportional to the number of synapses between the output and input cells. The proportionality constant depends on the specific pair of cell types and ROI, and determines how many synapses are made per unit of overlap.

The kernels and scaling factors were estimated by minimizing the error function

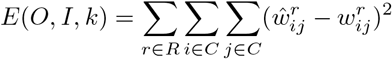

with 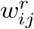 being the ground truth and the set *L* restricted to only the high-resolution columns.

To expand the resulting mean-grouped matrix to a full connectome matrix, the connection weight for each cell pair was taken to be the mean of the corresponding group multiplied by a noise factor. This noise factor was normally distributed with a global standard deviation parameter *σ* estimated by computing the standard deviation of the ROI connection weights divided by the corresponding noise-free extrapolated weights.

### Simulation and tuning

The circuit was modeled as a rate-based model with connectivity **W** based on the grouped connectome data. The firing rate dynamics were governed by

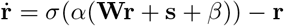

where **r** is the firing rate state vector, *σ* is a sigmoid activation function, *α* = 5 is a fixed gain parameter, *β* is a bias parameters, and **s** is external sensory input.

The weights were constructed as **W** = *E***W**_excitatory_ + *I***W**_inhibitory_ (where excitatory and inhibitory weights are normalized by their mean non-zero values), allowing us to separately tune the strength of inhibition and excitation, as well the bias *β*. These quantities are dimensionless and only their relative values are meaningful.

Exploring the parameter space (*E, I, β*) was done by identifying equilibria and following state trajectories from saddle points to stable points to trace out slow trajectories. The states on these trajectories were treated as candidate slow manifolds making up an approximate ring attractor.

Simulation was done in Python using forward Euler integration.

### Sensory input

The sensory input vector **s** consists of ring neuron inhibitory compass input, modeled as uniform inhibition of EPG neurons with local disinhibition based on their cosine alignment with the compass direction, and idiothetic GLNO rotational input that unilaterally inhibits one side of PEN cells, with strength proportional to the angular velocity.

### Identifying the ring manifold

The system’s equilibria **r**^*∗*^ were found by numerically solving 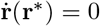 using initial guesses based on an ‘ideal’ bump at location *θ*, defined as

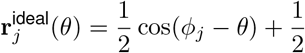

where *ϕ*_*j*_ is the preference angle of cells of column type *j* (assigned by their projections in the PB).

The states of EPG cells were pinned to their *A***r**^ideal^(*θ*) values for varying amplitudes *A* and locations *θ*, the system was relaxed, and the roots were found using scipy.optimize.root based on the resulting state as the initial guess.

Equilibria **r**^*∗*^ were classified as stable or unstable based on the eigenvalues of the Jacobian of the derivative 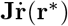. For an approximate ring attractor, we expect exactly one eigenvalue — whose eigenvector points along the tangent of the ring — to be close to zero, and all the rest to be strongly negative (in their real part). This ‘least stable’ direction corresponds to the direction in state space with the slowest dynamics. For all unstable equilibria, we thus perturbed the state slightly along this eigenvector and let the system relax to the adjacent stable state, tracing out its trajectory.

Additionally, PEN cells were unilaterally inhibited in order to trace out a cyclic trajectory corresponding to rotation of the bump. This was done by starting at an unstable equilibrium and simulating the system until a previous state is reached again, or for a fixed number of maximum steps, in which case such a cycle was determined to not exist.

### Model evaluation

States were projected onto the 2D plane using

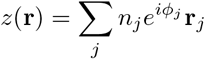

where *j* is the column type and *n*_*j*_ is the number of cells of that type.

This projection was then normalized as

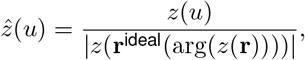

and the absolute value 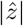 was used as the amplitude of the bump.

Using a grid search over (*E, I, β*), the model with the slowest maximum angular drift rate between equilibria (i.e. 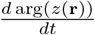 for states **r** along the ring manifold) along the ring was selected as the closest approximation of a ring attractor. This was done under the following constraints:

1. The projections of the states along the ring were not significantly biased in any direction, based on the mean value of their 2D projections (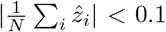, where *N* is the number of ring samples 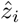).
2. Unilateral PEN inhibition resulted in a limit cycle in at least one direction depending on which side was inhibited.

### Simulated experiments

For the trajectory plots, the model state was initialized at a number of random bump-shaped locations in state space, i.e. at *A***r**^ideal^(*θ*) for random *A* and *θ*, and the system was then relaxed.

To determine the bump’s angular rotation speed in response to unilateral PEN inhibition, a range of levels of inhibitions were applied, and the system was simulated to look for a cyclic trajectory. The angular velocity response was computed as the mean angular velocity for the duration of each attempt to trace out such a trajectory.

For compass tracking simulations, agent headings were modeled as a moving average over a random walk. This heading was used as compass input to weaken the ER-like inhibitory EPG inputs. The rotation rate, in radians per time step, was multiplied by a factor of 15 (based on the fastest observed bump rotation speed) and used to unilaterally inhibit one side of PEN neurons. To remove external compass input, the weakening of ER-like EPG inputs was removed.

Virtual ablations were done by zeroing all outgoing connection weights from the ablated cell type.

## Data availability

All code is available at GitHub; all data will be made publicly accessible upon publication.

## End notes

## Acknowledgements

We acknowledge the facilities and scientific and technical assistance provided by the Microscopy Australia Facility at the Centre for Microscopy and Microanalysis (CMM), The University of Queensland. We are especially grateful to Nicole Schieber, Kathryn Green, Robyn Chapman, Rick Webb, and the late Roger Wepf for invaluable insights and access to the EM facility. We thank Robert Parton for collecting and providing live jack jumper ants. Worker honeybees were donated by Mandiyam Mahadeeswara and the Mandyam V. Srinivasan lab. We are grateful to Kevin Tedore (company: ‘Tedore Interactive’) for setup and maintenance of CATMAID and CAVE infrastructure and for providing ongoing technical support. We are grateful to Jan Funke for support with setting up our image segmentation pipeline, and to the CAVE team for continuous support with implementing and troubleshooting our CAVE deployment. M.E.S. was funded in part by an International Cotutelle Macquarie University Research Excellence Scholarship (iMQRES 2019060). A.N. was supported by the Australian Research Council Discovery Project grants, DP220102836 and DP250104339. S.E. was funded by Macquarie University international research training program (iRTP20225166). S.H. was supported by the European Research Council (ERC), under the Horizon 2020 and Horizon Europe frameworks (grant agreements 714599 – BrainInBrain and 01044220 – EvolvingCircuits), the European Innovation Council (EIC, grant agreement 101046790 — InsectNeuroNano), the Swedish Research Council (Vetenskapsrådet, grant agreement 2018–04851) and the Crafoord Foundation (grant agreement 20200709). Views and opinions expressed are however those of the author(s) only and do not necessarily reflect those of the European Union or the European Innovation Council. Neither the European Union nor the European Innovation Council can be held responsible for them. The funders had no role in study design, data collection and analysis, decision to publish, or preparation of the manuscript.

## Author Contributions

**Marcel E. Sayre** conceptualized the study, developed the imaging approach, acquired, processed and annotated the image volumes, analyzed and interpreted the data, developed and adapted analysis software, curated image and annotation data, designed figures, co-wrote the initial manuscript draft, and reviewed and edited the final version of the manuscript; **Valentin Gillet** developed and implemented image segmentation, proof-reading and analysis software, set up hardware environment for machine learning training, acquired all segmentation ground truth, annotated image data, contributed to data visualization, curation, data interpretation, manuscript writing and editing; **Nils Ceberg** developed and performed all computational modeling, analyzed, visualized and interpreted modeling results, contributed to figure design, writing and editing of manuscript; **Griffin Badalamente** developed, implemented and applied machine learning for automatic synapse detection, contributed to image analysis, annotation and interpretation, data curation, manuscript writing and editing; **Atticus Pinzon-Rodriguez** contributed to data annotation, curation, image analysis, training and supervision, review and editing of manuscript; **Nina Griggs** contributed to neuron annotation and proof reading, interpretation of data, manuscript review and editing; **Laia Serratosa Capdevila** contributed to neuron annotation and proof reading, interpretation of data, manuscript review and editing; **Ruairí Roberts** contributed to neuron annotation and proof reading, interpretation of data, manuscript review and editing; **Ebba S. Gunnarsson**: contributed to neuron annotation, interpretation of data, manuscript review and editing; **Felicia Szadaj** contributed to neuron annotation, manual synapse annotation, interpretation of data, manuscript review and editing; **Saroja Ellendula** contributed to neuron annotation and proof reading, interpretation of data, manuscript review and editing; **Anna Honkanen** acquired and processed samples, developed optimized fixation protocols, contributed to manuscript review and editing; **Ajay Narendra** supervised students and postdocs, acquired funding and resources, contributed to data interpretation, and manuscript writing and editing; **Stanley Heinze** conceptualized the study, acquired funding and resources, supervised students and postdocs, lead the overall project, contributed to data annotation, curation, analysis, validated and interpreted results, co-wrote the initial manuscript, contributed to figure design, lead writing and editing of final manuscript.

## Declaration of Interests

None to declare.

## Correspondence

Correspondence and requests for materials should be addressed to Stanley Heinze.

## Extended Data Figures

**Extended Data Figure 1:**
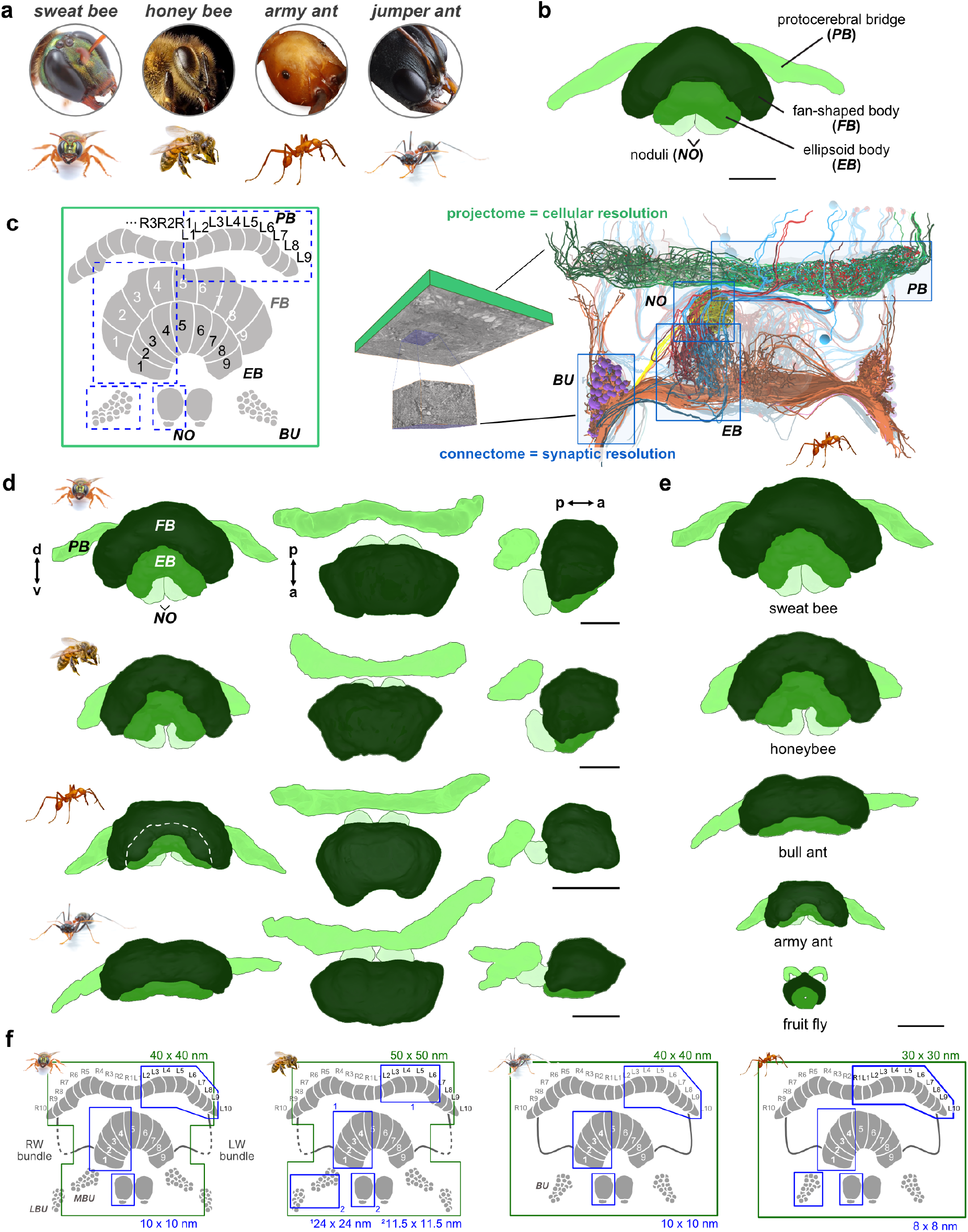
The central complex of all five species segmented from SBEM image data. **(a)** 3D reconstruction of neuropils in the sweat bee brain. Abbreviations: OL, optic lobe; MB, mushroom body; AOTU, anterior optic tubercle, AL, antennal lobe; CX, central complex; OC, ocelli; BU, bulb; d, dorsal; v, ventral; m, medial; l, lateral. **(b)** Central complex of the sweat bee (left) and fruit fly (right). **(c)** Central complex organization across species arranged by phylogeny. Icons denote similarities and differences relevant to navigational behavior. Divergence times (Ma) are from^76^. **(d)** Frontal, dorsal and lateral views of the central complex for each species (left to right). Species shown from top to bottom: sweat bee, honeybee, army ant and jumper ant. **(e)** Central complexes scaled for size comparison. **(f)** Schematic of SBEM image tiling, indicating overview (green) and high-resolution (blue) tiles and corresponding pixel sizes. All datasets were acquired at 50 nm z-resolution. Scale bars: 100 µm.

**Extended Data Figure 2:**
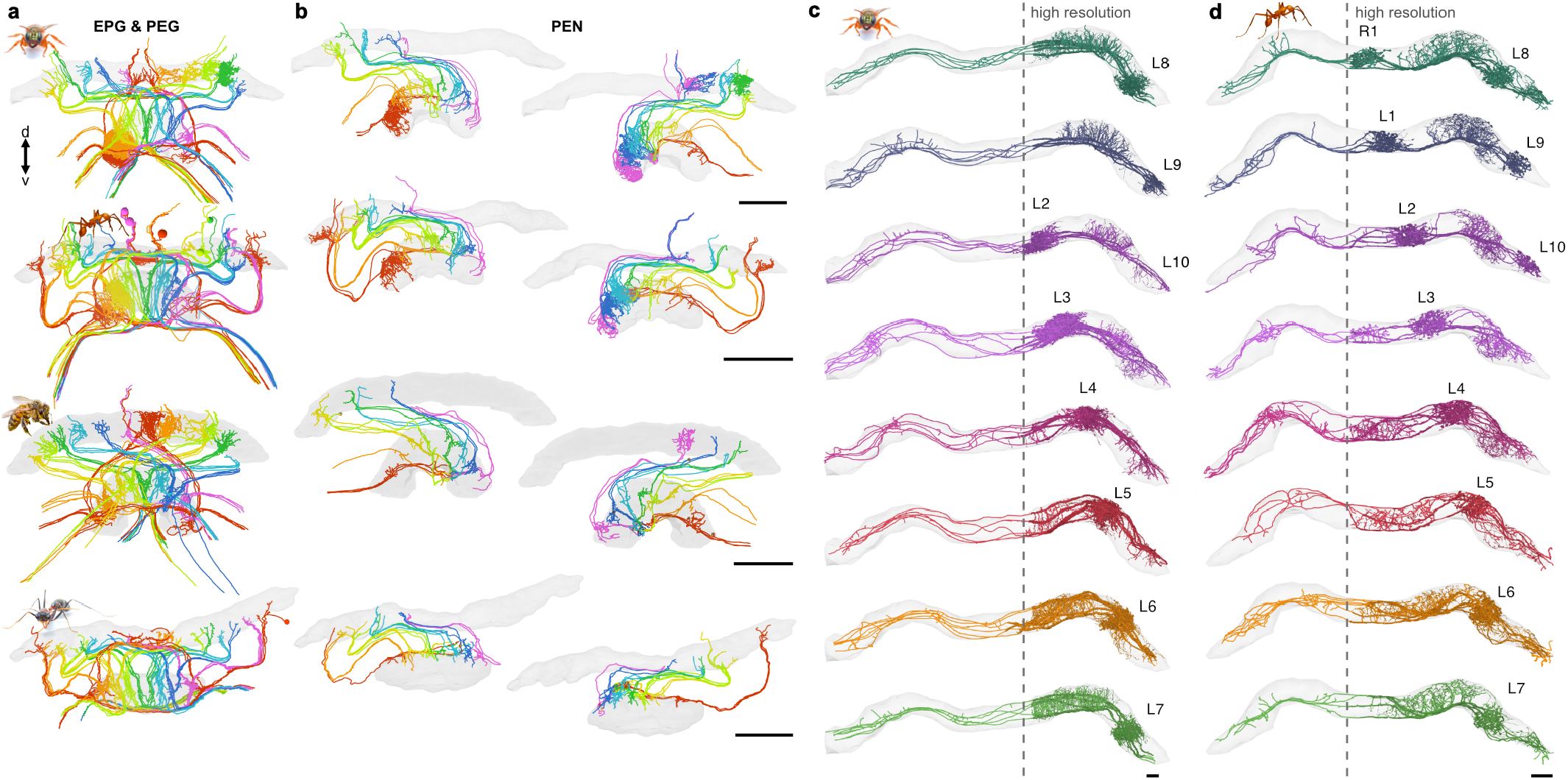
Projectome level reconstructions of neurons comprising the core head direction circuit. **(a)** Frontal views of manually reconstructed EPG and PEG neuron skeletons in ants and bees. **(b)** PEN neurons (combined PEN_a and PEN_b), shown separately for each hemisphere. Scale bars: 100 µm. **(c-d)** Frontal view of Δ7 cells in sweat bee (c) and army ant (b) protocerebral bridge. Dotted line indicates medial limits of synaptic-resolution image data. Scale bars: 20 µm.

**Extended Data Figure 3:**
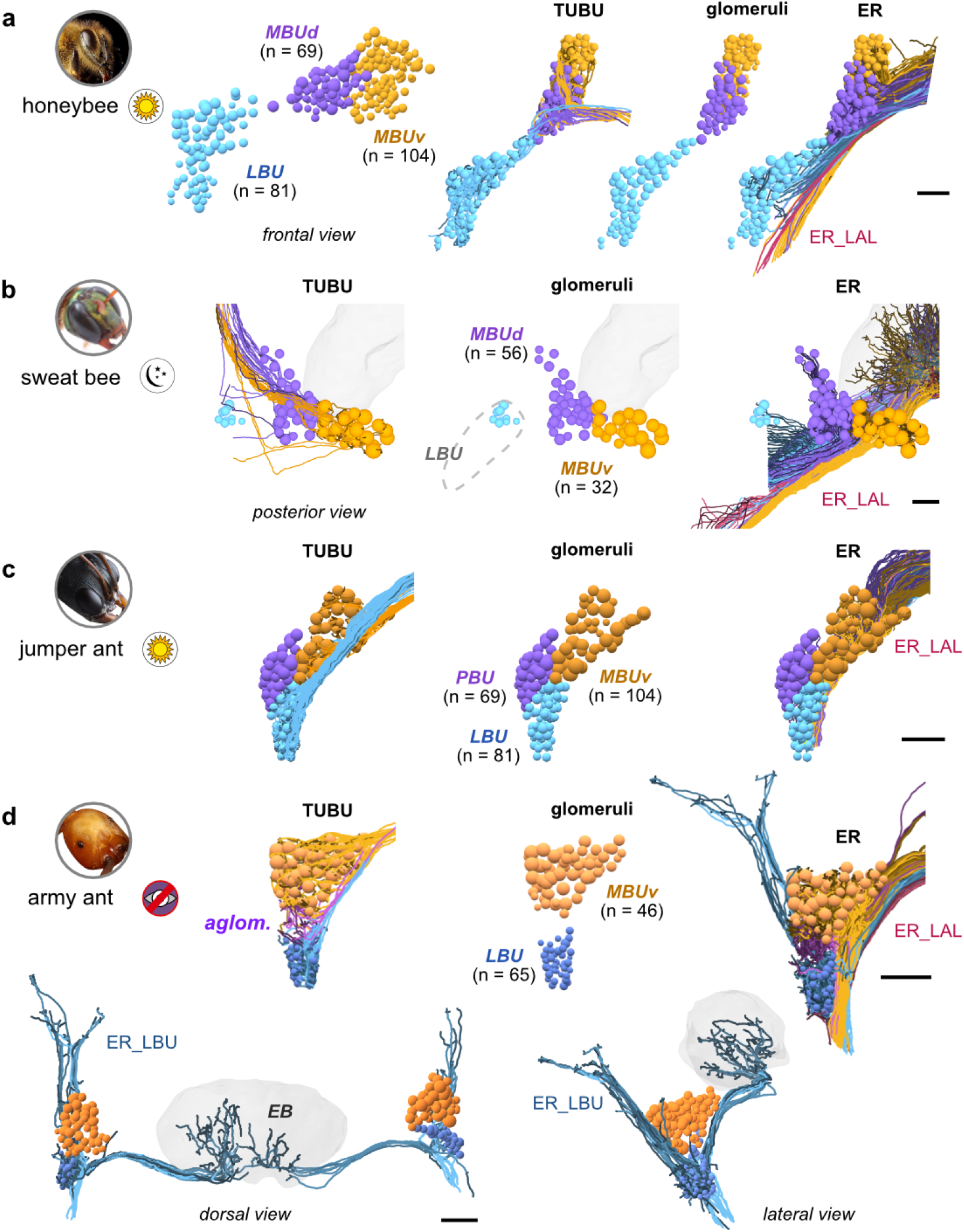
Glomerular organization of the bulb in the honeybee, sweat bee, jumper ant, and army ant. **(a)** Honeybee bulb subregions. (MBUd; dorsal medial bulb, MBUv; ventral medial bulb, LBU; lateral bulb). **(b)** Same as (a) for sweat bee. Grey dotted line indicates region not included in our image data. **(c)** Same as (a) for jumper ant. (PBU; posterior bulb) **(d)** Same as (a) for army ant. Also shown are a subtype of ER_LBU cells not found in the other species that possesses branches projecting toward the posterior protocerebrum. Scale bars: 25 µm.

**Extended Data Figure 4:**
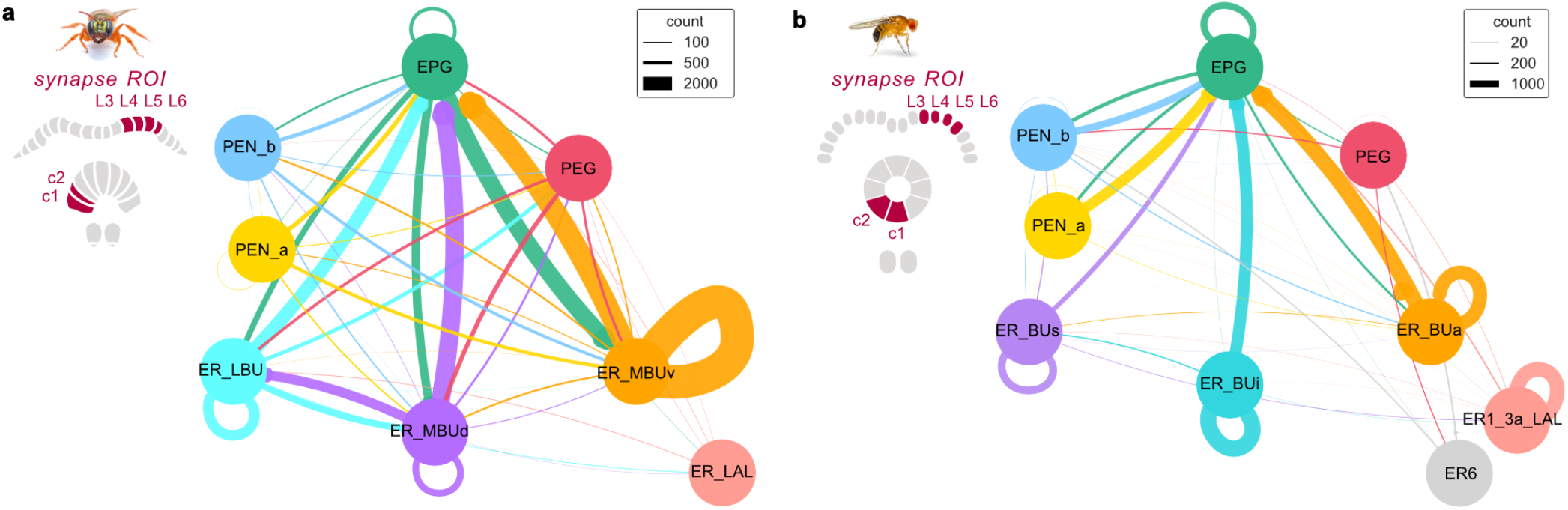
Connectivity of sensory input neurons in the sweat bee and fly ellipsoid body. **(a-b)** Type-level connectivity graphs of head direction neurons within the ellipsoid body of the sweat bee (a) and fly (b). To account for unproofread sweat bee ER neurons excluded from analysis, ER synapse counts were scaled by the total number of ERs of each type (see Methods).

**Extended Data Figure 5:**
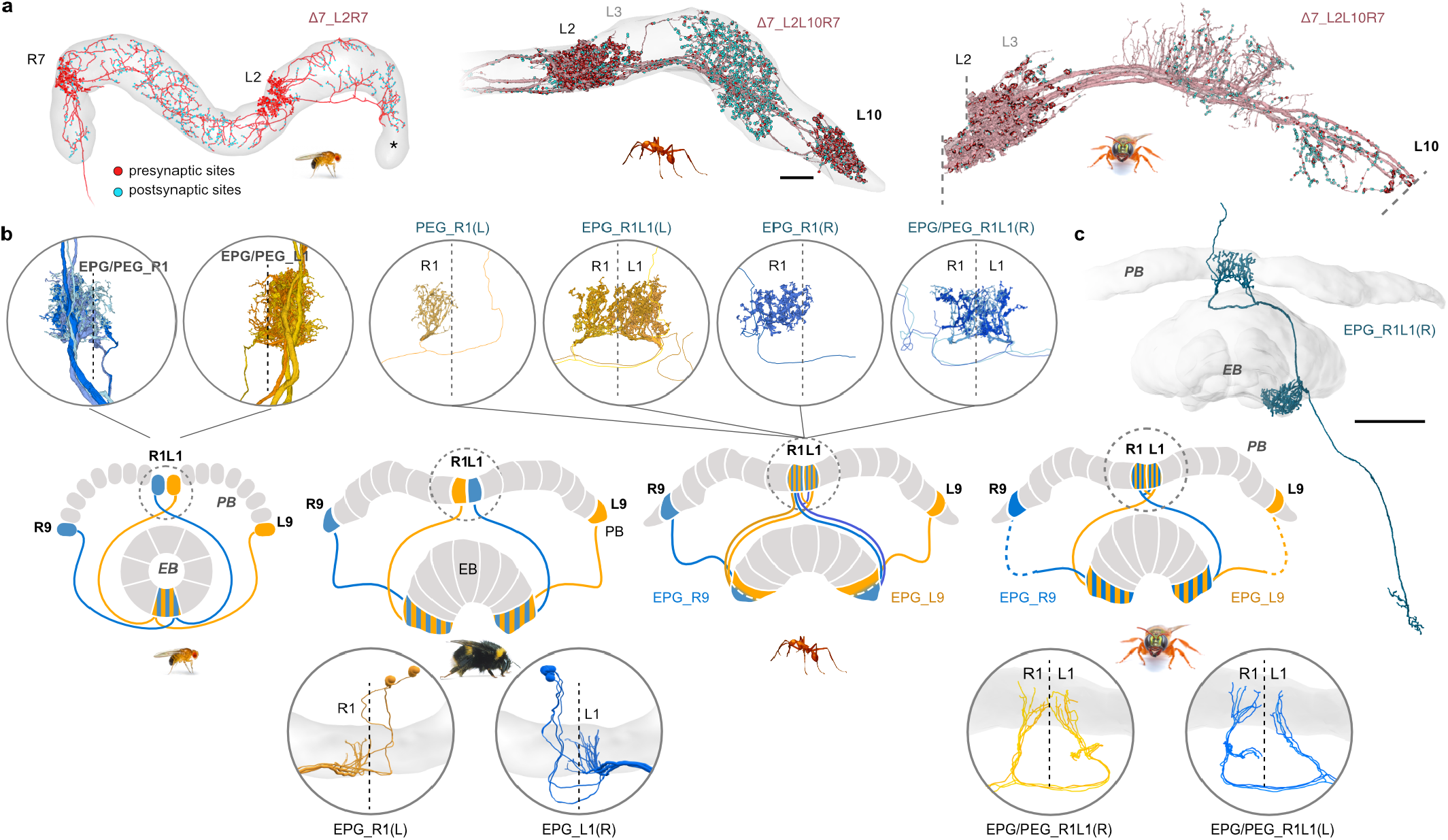
Species-specific specializations at the lateral extremes of the ring attractor. **(a)** Frontal views of fly Δ7_L2R7 and hymenopteran Δ7_L2L10R7 neurons, revealing a tenth output domain in the protocerebral bridge of the army ant and sweat bee. Dotted line in the sweat bee image indicates regions not captured in our image data (this includes the majority of the tenth PB column). **(b)** Projection patterns of EPG and PEG neurons at the innermost protocerebral bridge columns. Circular insets highlight branches of the individual midline EPG neuron types in the protocerebral bridge of *Drosophila* (left), bumblebee (second left), army ant (second right), and *Megalopta* (right). Bumblebee data from^37^. **(c)** Reconstruction of dye-filled single neuron of a midline EPG cell from *Megalopta genalis*. Scale bars: (a) 30 µm; (c) 100 µm.

**Extended Data Figure 6:**
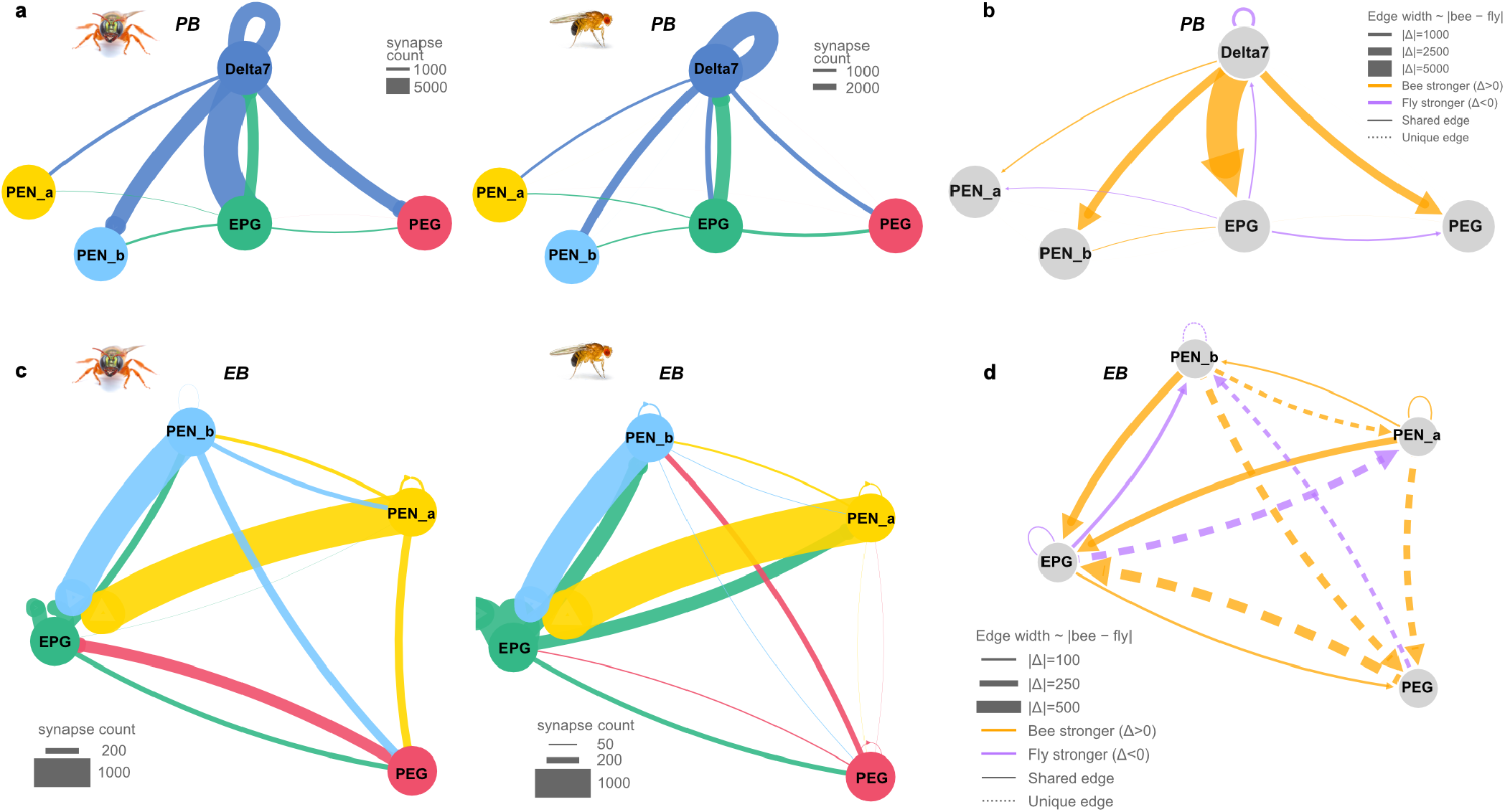
Type-level connectivity of ring attractor neurons in the sweat bee and fly. **(a)** Connectivity graphs of core head direction neurons in the protocerebral bridge of the sweat bee and fruit fly. **(b)** Difference graph highlighting species-specific connectivity among core excitatory neurons. Solid lines indicate stronger connections in bee (orange) or fly (purple); dotted lines denote connections unique to one species. **(c)** Connectivity graphs of core head direction neurons in the ellipsoid body of the sweat bee and fruit fly. **(d)** Same as (b) for neurons in the ellipsoid body.

**Extended Data Figure 7:**
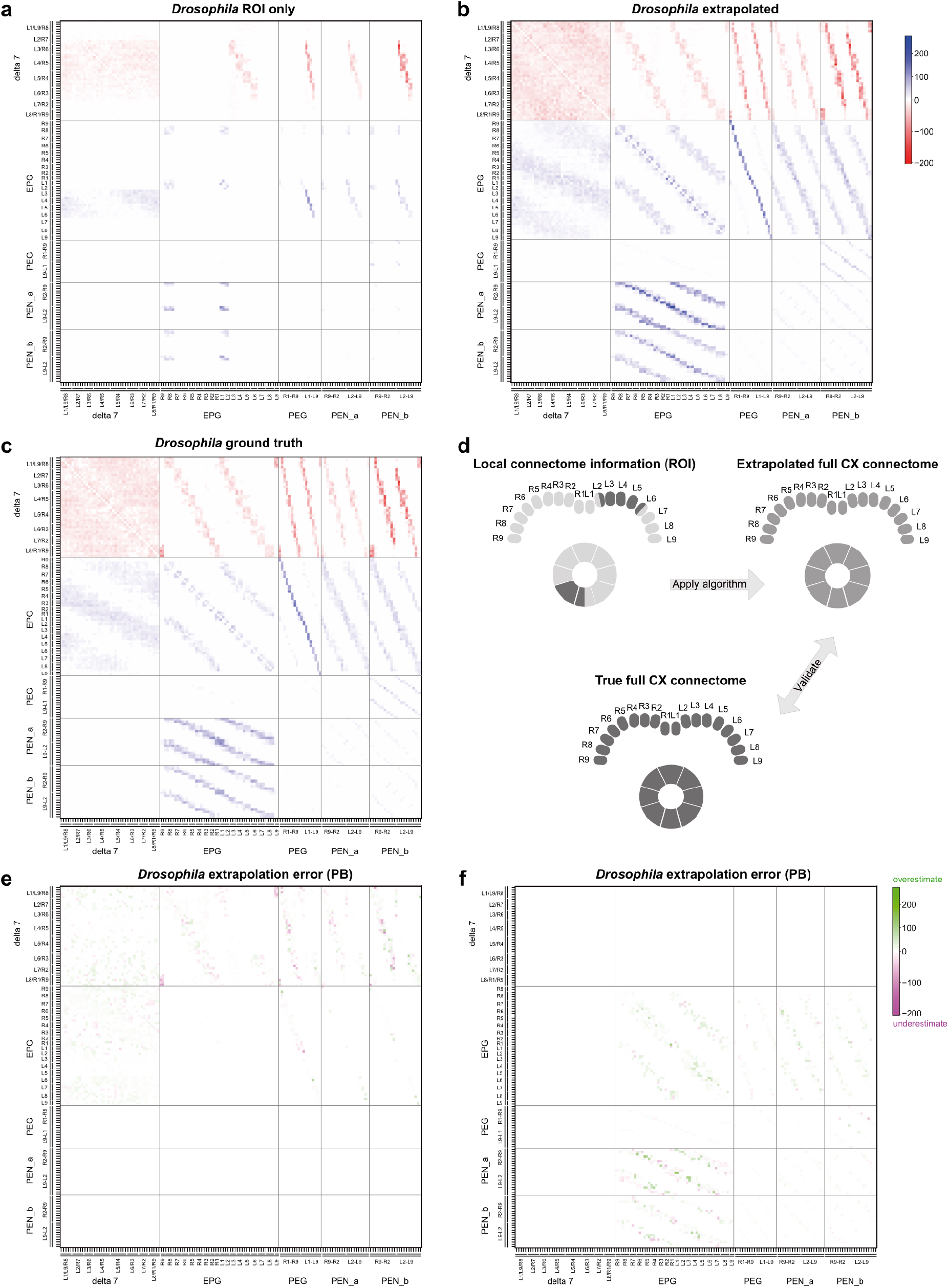
Validation of extrapolation method. **(a-d)** We applied the same extrapolation strategy as in the bee to data from *Drosophila*. Local connectome data were extracted from the fly connectome to match the regions of high-resolution data in the bee (a). The extrapolation strategy developed for the bee was then used to generate a full central-complex connectivity matrix (extrapolated matrix, (b)). This matrix was then compared to the ground-truth connectivity matrix directly obtained from the fly connectome (c). **(e**,**f)** Difference matrices between the extrapolated and ground-truth matrices of the fly, illustrating only minor, randomly distributed differences, likely due to noisy ground-truth connectome data. This demonstrates that valid, neuropil wide connectomes can be closely approximated using limited, local connectome data combined with low resolution projectome information. Grayed-out values were less than 2*σw*, where *w* is the noise-free extrapolated weight and *σ* is the standard deviation of the multiplicative noise added to the extrapolated connections.

**Extended Data Figure 8:**
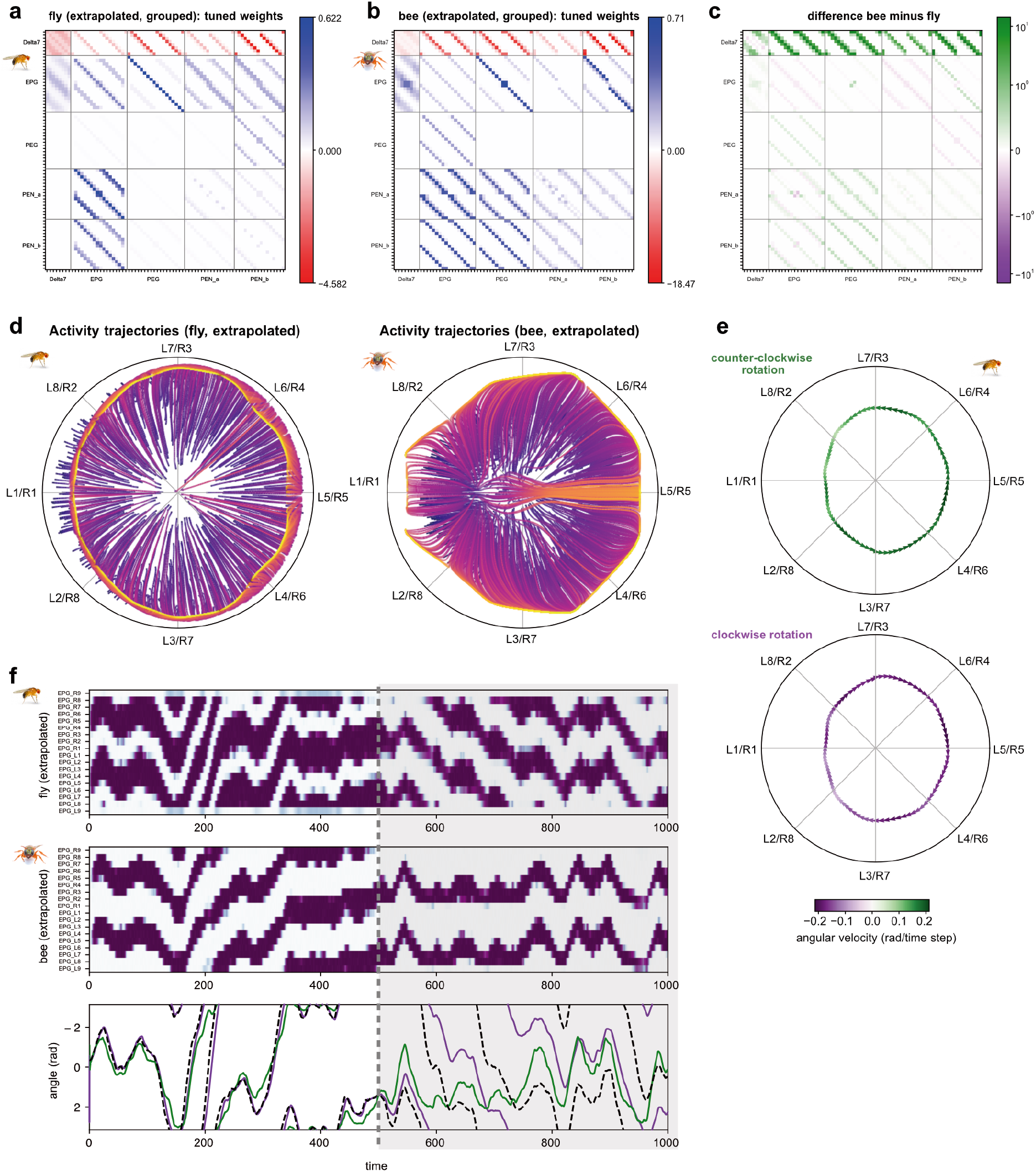
Comparison of bee and fly models. **(a**,**b)** Simplified connectivity matrices underlying the computational models for the fly and the bee, derived from extrapolated matrices by grouping of cell types within columns and by enforcing symmetry across the midline. Arborization kernels learned from local connectomes were used throughout the central complex. Shown are the normalized, tuned weights used for the models. **(c)** Difference matrix between bee and fly, highlighting the differences in neural connection weights in the tuned models. **(d)** Trajectories of neural activity states over time after initiating activity at random locations in state space. The ring-shaped structure demonstrates ring attractor dynamics in both species (left, fly; right, bee). The fly circuit more closely resembles an ideal ring attractor compared to the bee circuit. **(e)** Ring manifolds and drift speeds during imbalanced PEN neuron activation, mimicking rotational velocity input in the absence of visual input, shown for the fly-based model. More intense activation of PEN neurons in one hemisphere results in smooth movement of the activity bump either in clockwise or counter-clockwise direction along the ring manifold. **(f)** EPG neuron activity resulting from models based on either species tracks the heading of a simulated agent, both with and without compass input. The shaded area indicates time steps without allothetic input to the circuit, thus solely relying on idiothetic angular velocity inputs. Both circuits accumulate significant heading error without allothetic input, and in the bee model only traverses the midline in a narrow range of angular velocities.

**Extended Data Figure 9:**
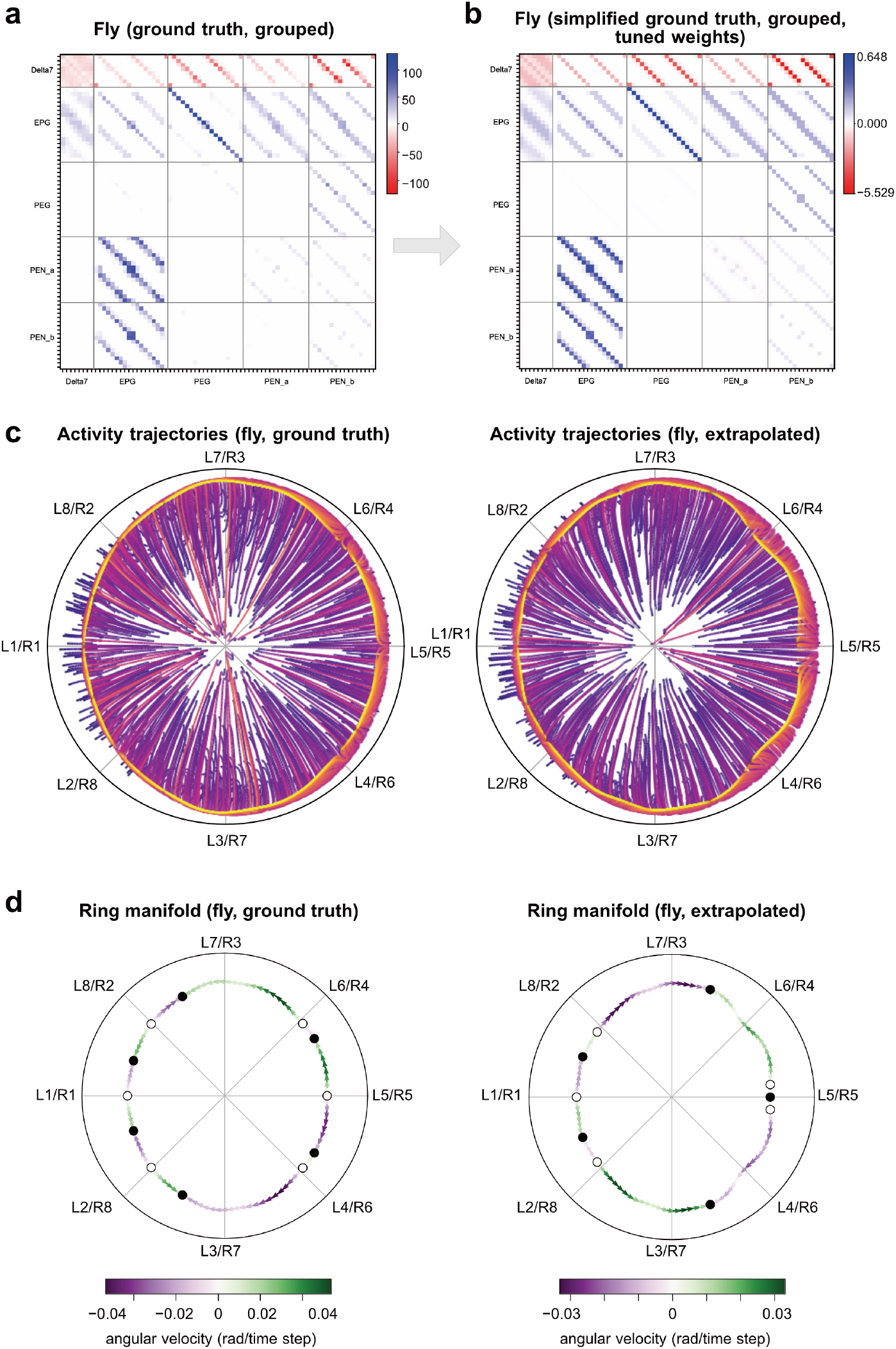
Comparison of models based on *Drosophila* ground-truth and extrapolated connectivity. **(a**,**b)** To validate the model based on extrapolation of local connectomics data in the fly, we have applied the same simplification procedure to the full connectome of the fly central complex. This included grouping of cells of the same cell types within each column as well as applying learned arborization kernels to yield initial connection weights. Rather than being based on partial data, here, these kernels were learned from the entire dataset, and thus represent the average arborization structure of the entire central complex, eliminating effects of local noise, asymmetries and inter-columnar variation. Shown is the grouped ground-truth connectivity matrix (synapse counts; (a)) and the simplified connectivity matrix (tuned weights, (b)). **(c)** Trajectories of neural activity states over time after initiating activity at random locations in state space. The ring-shaped structure demonstrates ring attractor dynamics. The dynamics of the extrapolated model and the ground-truth based model are near identical. **(d)** Ring manifolds for neural activity in the extrapolated and ground-truth based fly circuits, demonstrating highly similar ring attractor performance in either model. Circular axis: anatomical space in central-complex columnar coordinates. Shown are stable equilibria (filled circles), unstable equilibria (open circles), as well as local drift speeds of localized activity along the ring manifold. Slower drift speed indicates better approximation of an ideal ring attractor.

**Extended Data Figure 10:**
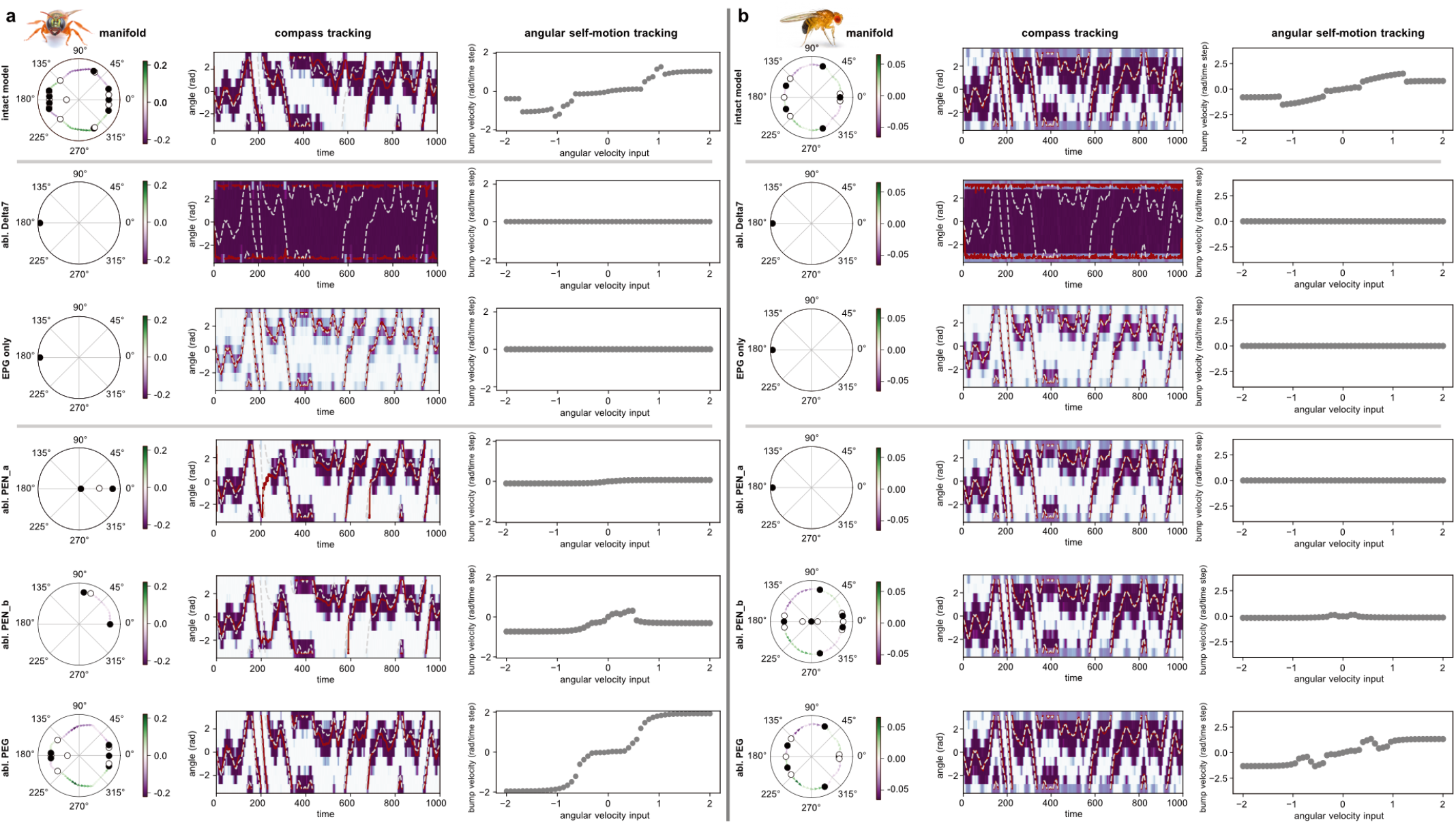
Ablation experiments in bee and extrapolated fly models. **(a)** Ring manifolds, EPG-neuron compass tracking ability, and self-motion based angular velocity encoding ability for different models of the bee circuit, in which different cell types were eliminated. Angular velocity input is modeled as asymmetric inhibition of all PEN neurons of one hemisphere (mimicking GLNO input to the noduli). Without Δ7 neurons, no localized activity bump exists in the central complex and activity spreads throughout the EPG population. With only EPG neurons present (all other cell types eliminated), simple visual inputs are directly encoded by EPG neurons, but no ring attractor dynamics exist. When PEN_a, PEN_b, and PEG neurons are eliminated, the dynamical properties of the system change. Both PEN subtypes are required for functional attractor dynamics and for angular velocity based lateral shifts of the EPG neuron bump. Ablation of PEG neurons had the least detrimental effect on the attractor dynamics, but resulted in much higher gain of the self-motion based angular velocity coding, i.e. a less stable activity bump. These simulation results align well with our finding that the PEG neuron connections are the most evolvable between flies and bees, and are thus suited to adjust detailed attractor dynamics in each species, while not interfering with core head direction coding. **(b)** As (a), but for the fly-based model (extrapolated model).

## Notes

### Competing Interest Statement

The authors have declared no competing interest.

### Summary of Updates

Figures slightly reformatted for clarity; Supplementary figures reformatted for clarity; figures S7,S8, S10 removed as only referred to once in the main text and partially redundant with the main figures; figures S2 and S3 combined into one figure; Figure captions updated and shortened for clarity; one author name corrected; figure cross reference errors fixed; author contribution section updated.

